# A Gene Regulatory Model of Cortical Neurogenesis

**DOI:** 10.1101/394734

**Authors:** Sabina S. Pfister, Andreas Hauri, Frederic Zubler, Gabriela Michel, Henry Kennedy, Colette Dehay, Rodney J. Douglas

**Affiliations:** Institute of Neuroinformatics, University of Zurich and ETH Zurich, 8057, Zurich, Switzerland; Department of Neurology, Bern University Hospital, 3010, Bern, Switzerland; University of Lyon, Université Claude Bernard Lyon 1, Inserm, Stem Cell and Brain Research Institute U1208, 69500 Bron, France

## Abstract

Sparse data describing mouse cortical neurogenesis were used to derive a model gene regulatory network (GRN) that is then able to control the quantitative cellular dynamics of the observed neurogenesis. Derivation of the network begins by estimating from the biological data a set of cell states and transition probabilities necessary to explain neurogenesis. We show that the stochastic transition between states can be implemented by the dynamics of a GRN comprising only 36 abstract genes. Finally, we demonstrate using detailed physical simulations of cell mitosis, and differentiation that this GRN is able to steer a population of neuroepithelial precursors through mitotic expansion and differentiation to form the quantitatively correct complex multicellular architectures of mouse cortical areas 3 and 6. We find that the same GRN is able to generate both areas though modulation of only one gene, suggesting that arealization of the cortical sheet may require only simple improvisations on a fundamental gene network. We conclude that even sparse phenotypic and cell lineage data can be used to infer fundamental properties of neurogenesis and its organization.

**Highlights:**

- Estimation of the cell states and transition probabilities of neurogenesis from experimental data.
- Design of an abstract gene regulatory network (GRN) whose dynamics implement cell states and their stochastic transitions.
- Detailed simulation of GRN-guided neurogenesis for mouse cortical areas 3 and 6.
- Different dynamics of neurogenesis of distinct cortical areas arise through modulation of only a single gene.

**In brief:** Pfister et al. show how sparse phenotypic and cell lineage data can be used to infer a small abstract gene regulatory network (GRN), which, when inserted into model precursor cells, is able to control in a distributed manner the quantitative cellular dynamics of neocortical neurogenesis.

## 3. Introduction

Unlike human engineered systems that are explicitly designed and constructed, the rules for self-construction of biological organisms are implicit in the information contained in their initial cells. Although many details of this remarkable process have been described experimentally, there are as yet no detailed generative models that describe formally the principles of control and global coherence amongst proliferating, locally independent, cellular agents. Here we describe a number of significant advances toward this goal in the context of the development of the laminated neocortex from its neuroepithelial precursors. We show how sparse phenotypic and cell lineage data can be used to infer a small abstract gene network, which, when inserted into model precursor cells, is able to steer in a distributed manner the quantitative cellular dynamics of neocortical neurogenesis. Our results offer an insight into principles of physical self-construction of biological neural networks.

Neocortical pyramidal cells are generated, and migrate to form a type specific lamination, however, the cellular mechanisms that underly this cortical neurogenesis remain elusive (Greig et al., 2013). Cortical neurogenesis begins from a sheet of neuroepithelial stem cells. These cells differentiate predominantly into radial glial cells (RGC) (Hartfuss et al., 2001; Miyata et al., 2001; Noctor et al., 2001, 2002; Anthony et al., 2004). RGCs divide at the apical surface of the ventricular zone (VZ), where they undergo stereotypical sequences of cell divisions: Symmetric divisions lead to similar offspring and amplify the pools of precursor cells; asymmetric divisions give rise either to various intermediate precursors, (Franco and Müller, 2013; Guo et al., 2013), or directly to cortical neurons (Heins et al., 2002; Malatesta et al., 2003; Anthony et al., 2004; Cárdenas et al., 2018) (reviewed in Götz and Huttner (2005)). Some precursors are restricted to the VZ (Haubensak et al., 2004; Miyata et al., 2004; Noctor et al., 2004), and are the major source of the deep layer pyramidal neurons. Other precursors form a second germinal layer, the subventricular zone (SVZ). There they undergo a few rounds of symmetric division and generate neurons largely fated for the superficial layers (Noctor et al., 2004; Kowalczyk et al., 2009).

The genealogical lineages whereby the neuroepithelial stem cells give rise to differentiated neurons are only partially known (Haydar et al., 2003; Noctor et al., 2004; Gao et al., 2014; Vasistha et al., 2015; Telley et al., 2016; Beattie and Hippenmeyer, 2017; Kaplan et al., 2017; Zhong et al., 2018). Every cell in the lineage has the same genotype, but the phenotype of each cell is due to its particular gene expression pattern, and interaction with environmental factors. The lineage tree describes the genealogy and division history of successive precursors, where each cell is associated with a particular phenotype. Ideally, the structure of the lineage tree should reflect the progressive restriction of cell fate. It would exhibit the variety of successive precursors that could be generated as neurogenesis proceeds, and thereby offers insights into the mechanisms that lead to the generation of experimentally observed neural cell types.

Although recent work points to an orderly and deterministic proliferation, and neurogenic behavior of precursors (Gao et al., 2014), the underlying organization of their lineage trees are not completely known. In principle, the progression of cell types through the tree can be characterized by their phenotypic description. The overall phenotype of a given cell can be represented as a vector of features *f* = {*f*_1_, *f*_2_, *…, f_n_*} that include its gene expression pattern, morphology, biochemical or physiological properties, and behavior. Some of these features may be observable, but others are hidden. We assume that this vector of cell features is conditioned by the internal unobservable cell state *S* that completely explains their distribution. The individual genealogical trees are the result of particular cell states, and the probabilistic transitions between them. Thus, the process of neurogenesis can be described in two complementary ways: The Cell Lineage Tree (CLT) that describes the genealogical relationship between the individual cells generated during development; and the State Diagram (SD) that describes the possible states that cells may take, and the stochastic transitions between these states. The functional mechanism underlying these descriptions is the mitotic process and its interaction with the gene regulatory network (GRN). Our challenge is to estimate the distribution of CLTs; to identify their underlying states and transitions; and then to posit a biologically plausible generative mechanism for their occurrence.

The purpose of this paper is to show that even sparse phenotypic and cell lineage data can be used to infer fundamental properties of neurogenesis and its organization. We begin by using previously published data to derive a stochastic state transition model of cortical neurogenesis, and from this we implement an abstract gene network that carries out the stochastic process. We then use a simulation of physical cell growth and mitosis to demonstrate that this GRN is able to steer in a distributed manner the quantitative cellular dynamics of neocortical neurogenesis.

## 4. Results

### 4.1 Cell lineage Trees

The Cell Lineage Tree is an acyclic directed graph in the form of a rooted binary tree, in which the vertices represent physical cell instances, and the directed edges represent the genealogical relationships between mothers and their daughter cells. The root of the tree is the earliest stem cell (neuroepithelial cells in this case); the internal nodes of the tree are dividing multipotent or pluripotent precursor cells; and its leaf nodes are non-dividing terminally differentiated cells (neurons and glial cells).

Measurements of lineage subtrees indicate that at least in vertebrates the lineage mechanism is stochastic rather than deterministic (He et al., 2012). Thus, vertebrate lineage trees form a distribution over possible genealogies. When two new cell instances are generated by mitosis, fate transitions occur between the precursor and its offspring. If the precursor divides symmetrically it will produce two daughters with identical cell fates, and thus identical phenotypes. However, if it divides asymmetrically, the precursor will produce two cells that inherit distinct gene expression products, and as a consequence may have different cell fates. In principle, we could measure the feature vector *f* over all cell instances. But such an exhaustive description is not yet technically feasible. Thus, for the present purposes, we assume that the feature vectors can be observed only over terminally differentiated cells. That is, we can observe and classify the phenotypes of terminal cells in terms of their neuronal morphology and behavior. Figure 2A shows a simple CLT, for purpose of explanation. The terminal states of this CLT are categorized into three types (*A, B, C*) based on a set of features {*f*_*A*_, *f*_*B*_, *f*_*C*_}, which we assume can be observed only in terminal cells.

### 4.2 Cell Lineage Trees for mouse cortical neurogenesis

We obtained estimates of the distributions of terminal neuronal types in mouse area 3 and 6 from the work of Polleux et al. (1997a), who used pulse ^3^ *H*-thymidine injections made throughout corticogenesis to measure the variation of cell cycle duration, cell cycle exit probability and laminar fate as functions of developmental time. Following their data and methods, we computed the temporal generation of neuronal types by numerical solution of the continuous differential equations describing cell proliferation and differentiation (Polleux et al., 1997b) (Figure 1). We then used these population distributions together with a probability-generating function (Bremaud, 1988) to generate probabilistically instances of cortical cell lineages (Figure 1).

**Figure 1.**
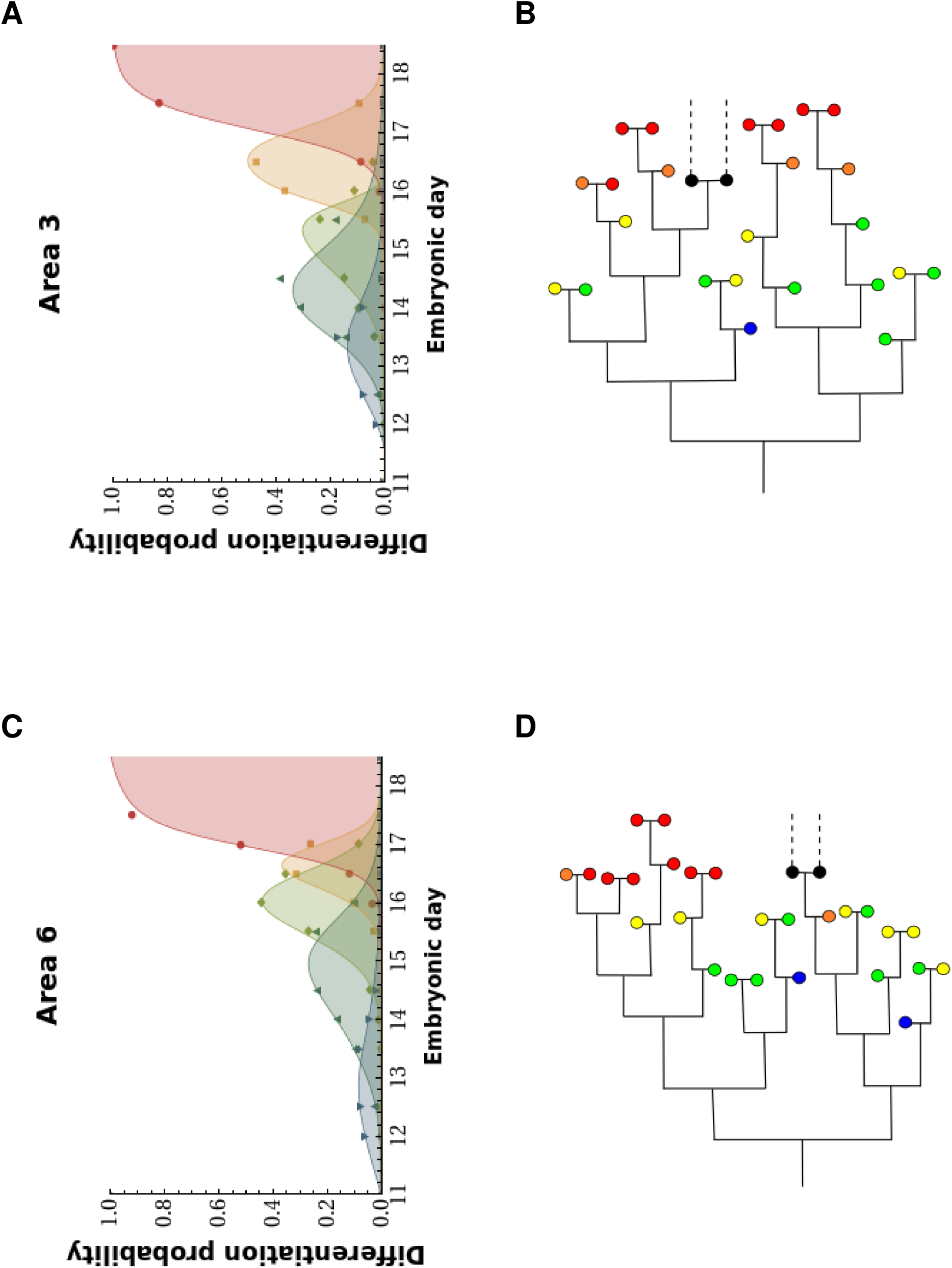
Probabilistic generation of lineage trees. Lineage trees are generated by sampling from the experimentally determined probability distribution (re-analysed from data of Polleux et al. (Polleux et al., 1997a)). (**A,C**) Probability distributions for area 3 and 6. Points, experimental data; lines, fits to data. (**B,D**) Example of sampled lineage trees. Trees layed out to correspond with the time axis of the experimental data. Black: precursor cell; blue: layer 6b; green: layer 6a; yellow: layer 5; orange: layer 4; red: layer 2-3; dashed lines, proliferation of glial precursor cells (not modeled).

**Figure 2.**
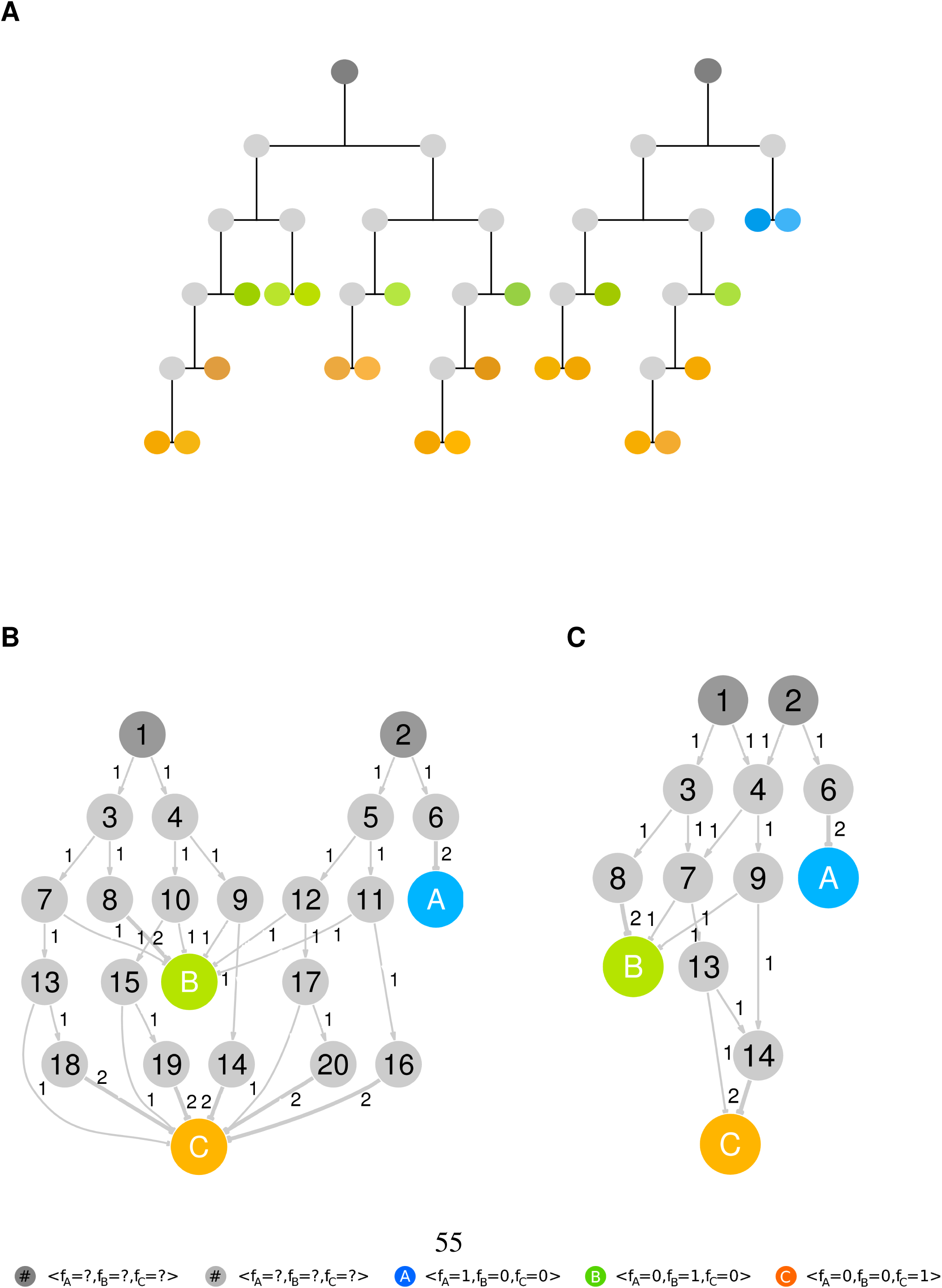
Cell Lineage Trees and their corresponding State Diagram. (**A**) Illustrative example of two cell lineage trees. Each node corresponds to a cell, and connecting edges to cell divisions. Two progenitor cells (dark gray) divide to form various hidden proliferative cells (light gray) and thereby give rise to 22 observable, terminally differentiated cells. Colors represent vectors of observed features (*f*_*A*_, *f*_*B*_, *f*_*C*_). (**B**) State Diagram describes how the various cell states in lineage trees of A) are related. The hidden states are numbered in correspondence with each hidden cell in the lineages. Colored cells in the lineages have the same phenotypic features and so are represented by only a single state here. Edges between nodes indicate the transition probabilities *p*_*ij*_ from states *i* to *j* (the probabilities account for 2 offsprings per division). (**C**) Reduced State Diagram obtained by combining the redundant hidden states of B).

### 4.3 State Diagrams

An alternative view of neurogenesis is one that describes the underlying generic cell states and their transitions, rather than the genealogical relationships between particular cell instances. We will call this alternative view the State Diagram (SD). It is a weighted directed graph whose vertices represent cell states, and whose weighted edges represent the stochastic transitions between states that occur at cell mitosis. Whereas the CLT describes both terminal cell identities and their individual ontogenies, the SD explains the experimentally observed numbers and dynamics of production of neuronal types in terms of state transition probabilities.

The SD begins from an initial precursor cell state; for example, the state of a neuroepithelial cell. When a cell undergoes mitosis, it generates two daughter states that will themselves generate subtrees of states, until a terminal state is reached. Because the SD vertices are states and not specific cells, cells that have exactly the same state are represented by the same single vertex. The numbers of cell transitions between one state and a different one are accounted for in the probabilistic weights of the edges that join the states. However, the sum of the probabilities across all the possible transitions away from a mother state is 2 not 1, because always two daughter states must be generated.

The SD can have different degrees of resolution, according to the mapping of individual physical cells to their possible underlying cell states. Trivially, any collection of lineage trees can be encoded exhaustively by an SD in which each and every cell instance is assigned to its own unique state (Figure 2B). Although a high resolution representation of this type is easy to generate, the number of states increases exponentially with the complexity of the cell lineage trees. The SD soon becomes intractably large, and the number of unique states and transitions rapidly exceeds the amount genetic information available to encode it.

A more suitable mapping of cells onto states assumes that biological processes are often best explained by models with low but noisy dimensionality. This is likely true for cell lineages, where only a very small set of all possible internal genetic expression profiles are visited by cells during development (Kauffman and Kauffman, 1993), and because very similar cell division sequences occur across the distribution of all lineage trees. Such a reduced encoding involves collapsing high dimensional graphs into subgraphs that have the same or similar underlying states and transitions. The example SD (Figure 2C) shows the principle of this reduction of redundant subtrees. The result is a more compact representation that describes the same developmental process, but using fewer states.

The general problem is to find such a low dimensional SD that is still able to account for most of the variance in the experimental data. We approached this problem by spectral clustering (Chung, 1997; von Luxburg, 2007), a type of clustering algorithm that can be applied to graphs. Our goal was to obtain an appropriate embedding of the full dimensional SD into a similarity matrix, such that the pairwise distance between cell states in the embedding space reflects their similarities in terms of terminal cell types than those two states give rise to. Once the full SD is embedded into an Euclidean space, simple algorithms such as hierarchical clustering can be used to cluster cell states into smaller subsets and thereby generate a lower dimensional, more easily interpretable SD representation of the cell lineage.

Since the SD states can be characterized by feature vectors, the reduced SD also models implicitly the statistical distributions over the feature profiles characteristic of each state, and the genealogical relationships between these feature states. Unfortunately we do not have data for the internal nodes of the SD (but see (Pfeiffer et al., 2016)). However, the feature vectors for the terminal states are known, and so we can estimate the feature profiles of the hidden vertices by propagating the known features backward into the hidden network. In this way the precursor states are mapped to corresponding linear combinations of terminal features. These profiles are a prediction of the contributions of the various precursors to the different final neuronal fates. For convenience we visualize these relationships by suitable coloring of the SD graph. The feature vectors of terminal states are associated with unique color vectors. These colors are then propagated backward into the network as proxies for features. The ‘colors’ of the precursor cells provide a visual impression of the fates to which they will contribute (Figure S2 and Figure S4). The SD states are an estimate of the hidden biological cell states *S*. For example, we may take this estimate to be *f*. And so each node of the SD is labeled with a vector whose elements correspond to experimentally observable features *f*_*j*_, such as the expression of a particular set of genes, or morphological features.

### 4.4 State Diagrams for mouse cortical neurogenesis

We used our spectral clustering method to estimate the SD underlying the development of cortical areas 3 and 6 of the mouse. The dynamics of cellular division and differentiation during development of these areas have been quantified using the mitotic history technique, which selectively monitors the proliferative behavior of defined cohorts of precursor cells generated at particular time points (Polleux et al., 1997b; Dehay and Kennedy, 2007). However, the behavior of the individual lineage trees supporting these population dynamics is unknown. There-fore we reconstructed probable lineage trees by sampling from the experimentally determined cell distributions (Figure 1). While the topologies of these trees are stochastic, their overall distribution is constrained by the experimentally observed distribution over different terminal cell fates.

We analyzed 60 such reconstructed lineages from area 3 and 6 of the mouse cortex. These lineages contained a total of 3263 cell instances (1549 in area 3 and 1714 in area 6). The terminal cells were labeled as either *Layer 6b* (L6b), *Layer 6a* (L6a), *Layer 5* (L5), *Layer 4* (L4), *Layer 2*/*3* (L2/3), or *Glia*. Precursor cells were labeled as *Unknown*. The complete, unreduced, SD was composed of 6 terminal states; with 765 unknown precursor states in area 3 and a further 848 unknown precursor states in area 6. Spectral clustering for both areas was performed on the combined dataset. The combination of data allows the method to exploit possible similarities between the SDs of the two areas (Figure 3).

**Figure 3.**
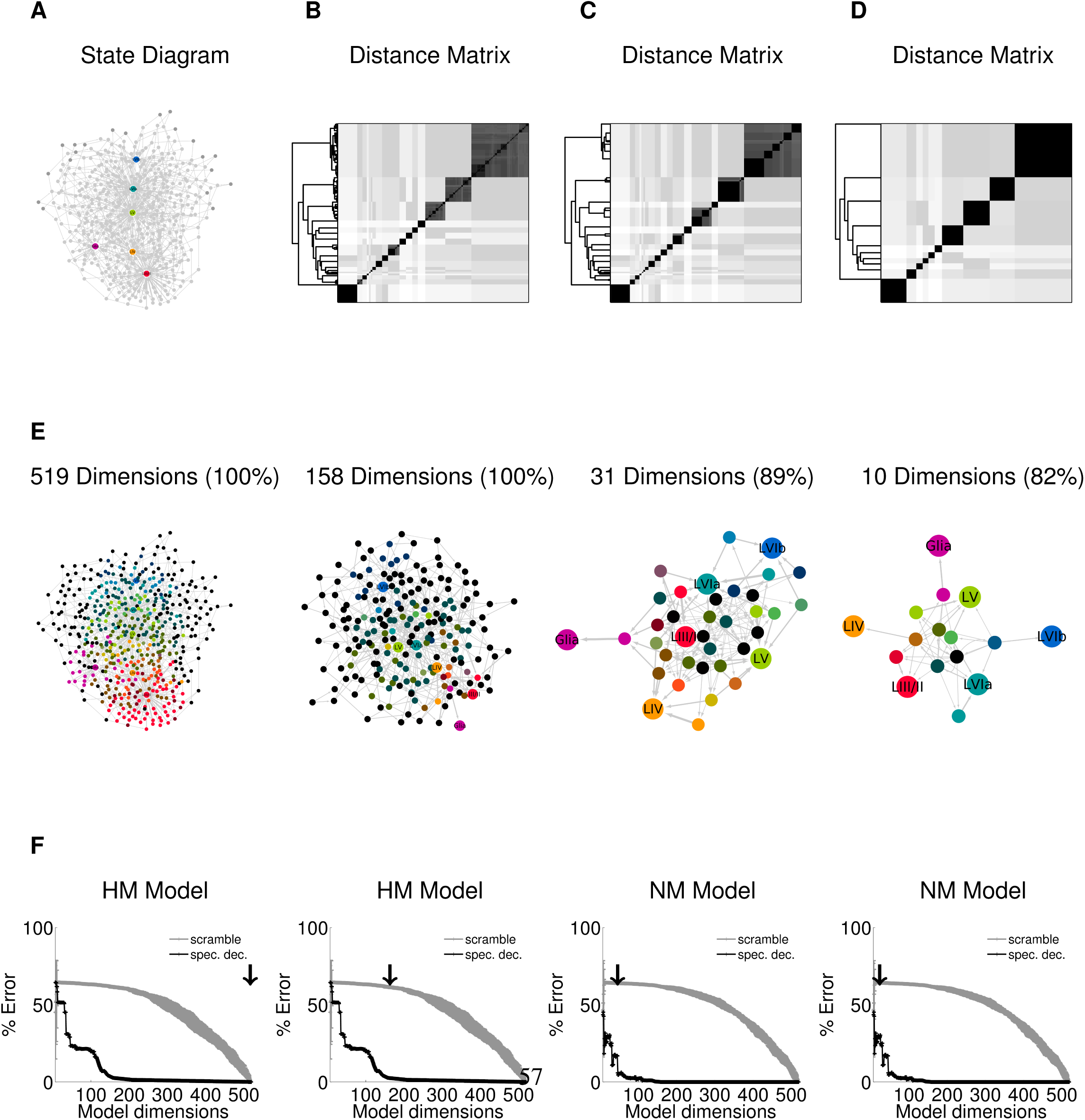
State Diagram of cortical area 3 and 6. (**A**) State diagram of cortical lineages in area 3 and 6 combined. Nodes represent cell states, arrows state transition probabilities. Cell states are labeled: blue: layer 6b; green: layer 6a; yellow: layer 5; orange: layer 4; red: layer 2-3; glia: pink, unknown; gray. Initial states are depicted as dark gray. (**B**-**D**) State clustergrams of computed distance between every state pair with dimensions *D* = 519, *D* = 158, *D* = 31, and *D* = 10 (percentage of data represented in parenthesis). Dendrograms indicate hierarchical binary linkage of states. (**E**) Spectral label propagation on models, where each nodes is colored according to the estimated feature distribution. (**F**) Model error as percentage of the correct final cell states distribution for spectral clustering (black) versus random model (gray, standard deviations on 100 trials). HM, Homogeneous Markov model; NM, Non-Homogenous Markov Model. Black arrow indicates dimensionality of model.

The original data is fully described by a SD of 519 dimensions, in which each cell has a corresponding state. Similar states generate cells with identical fates, and so can be collapsed into a unique state leading to a reduced SD with only 10 dimensions with negligible loss of accuracy. Models with even fewer dimensions are also able to describe the data, but with less accuracy. In order to compare the performance of SD models of different dimensions, we estimated the model error as the number of incorrectly generated terminal cells types over the total number of cells produced at the end of the developmental process. This error was compared against that of a complementary scrambled model, obtained by random permutation of cell states.

The accuracy of the SD models for area 3 and 6 was assessed for the homogeneous (HM), the non-homogeneous (NM) and the time-dependent (TM) Markov process. In the HM model, transition probabilities are independent of time, and so at low model dimensions the cell output distributions have long tails because of small state transition probabilities, which cause a small proportion of cells to undergo many rounds of division (Figure S6 and S7). Convergence to the target distribution occurs only after a great number of cell divisions, which is unrealistic for biological processes. We therefore introduced time dependence by applying age-dependent probability distributions in the NM model: Each state has unique outgoing transition probabilities, and a maximal number of possible self-replicative divisions. This assumption truncates the long tails of the HM approach, forcing cells to progress through the differentiation path. Finally, in the TM model, each transition probability is computed for each round of cell division. This model reproduces accurately the cell distributions as well as their temporal dynamics. However, this accuracy comes at the cost of a large number of parameters. By contrast, the HM model requires a large number of cell states for an accurate prediction. Both cortical areas are best described by the NM model, which is able to reproduce closely the system dynamics, and offers a good trade-off between model complexity (31 or 10 dimensions) and model accuracy (11% or 18% model error) (Figure 4A, B).

**Figure 4.**
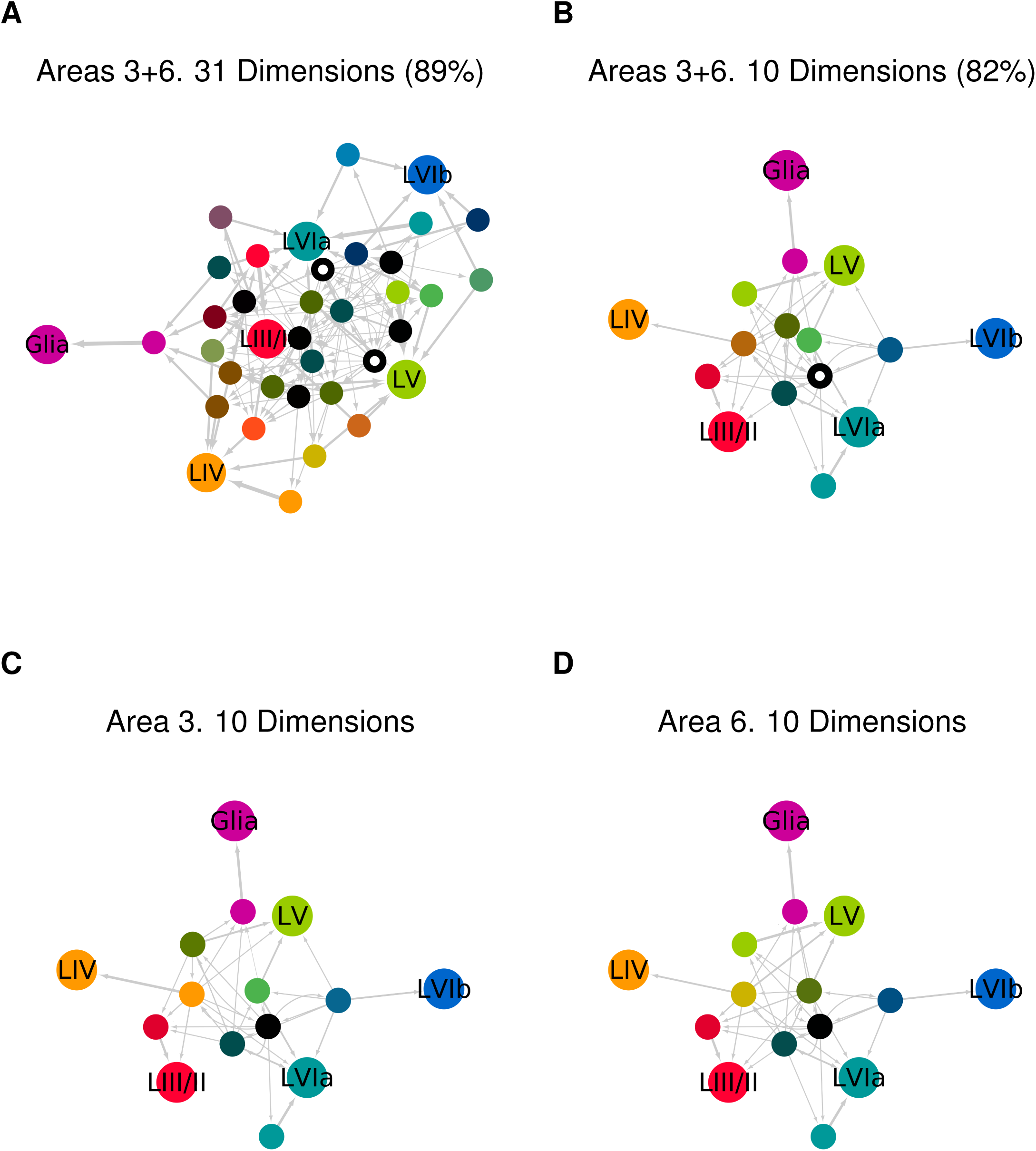
State Diagram details. (**A**-**B**) State Diagrams describing the combined lineages of areas 3 and 6. These 31 and 10 dimensional diagrams are enlarged from Figure 3. The initial precursor population(s) in these two cases are marked by centered white dots. The 31 dimensional SD declares a small second precursor population, whereas the 10 dimensional case collapses these two into a single initial population (with a small loss in ability to capture the experimental data). (**C**-**D**) Comparison of the two reduced State Diagrams for areas 3 and 6 respectively. The subtle differences can be seen in the shades of the three green/ocre small nodes in the upper left quadrants of the networks. The differences in shade indicate slight differences in predisposition towards terminal fates. (Networks enlarged from Suppl.Figures S6 and S7).

The NM 10 dimensional SD model explains 82% of the data, and is the most visually intuitive for reasoning over the logic underlying the developmental processes of area 3 and 6. The black node (with centered white dot) represents an initial homogeneous population of precursor cells, which then divide into subpopulations of precursor cells having different neurogenic potentials. A small proportion of cells are fated very early on to develop exclusively toward granular (L4) or supragranular layers (L2/3); and a large pool of heterogeneous precursor cells are less fate restricted (Figure 4B). The 31 dimension SD model is more precise: It explains 89% of the data, but it is less intuitive. A striking difference of this model with respect to the 10 dimension SD case, is the presence of two distinct initial populations that develop differently according to their fate restriction (Figure 4A). It is noteworthy that the precursor pool has some degree of plasticity in the sense that many cell states have bidirectional transitions, as has been observed in the cortical lineages of primates (Betizeau et al., 2013).

The SD’s above were computed over the combined lineage datasets for areas 3 and 6. However, we track the contributions of each dataset, and so it is straightforward to decompose the combined SD into the separate SDs describing each area (Figure S5). The reduced SDs for area 3 and 6 are strikingly similar (Figure 4C, D), suggesting that only minimal changes in a single model are sufficient to explain observed differences of neurogenesis in individual areas.

### 4.5 Estimates of SD gene expression patterns

So far we have interpreted the SD in terms of its propagation of terminal cell fates that are largely morphological, e.g. L2/3 pyramidal cell. However, SD models can also be interpreted in the light of the underlying gene expression process. For example, one might choose for features {*f*_1_, *f*_2_, *…, f_n_*} the real, observed transcription factor expression levels. Such data were not available to us at the beginning of this project. However, for illustration of the principle we used calibrated gene expression levels in cortical neurons obtained from a transcriptome atlas of cortical layers in the adult mouse area 3 (Belgard et al., 2011). Of the 11411 gene probes used in that atlas, we consider only the subset of 1751 transcription factors. We applied *k-means* clustering to this dataset and thereby identified 12 clusters of transcription factors that have similar expression patterns across the cortical laminae (Table S1). Each lamina is associated with one of the terminal neuronal types, and so each neuronal type is associated with a characteristic distribution across the 12 transcription factor clusters. Because the clustering is based on adult expression data, the distributions of the feature vectors are known only for terminal cell fates. However, as described above, our spectral clustering method can be used to propagate the adult values backward into the lineages and thereby provide a prediction of the expected transcription factor profiles to be found in the various SD precursor states (Figure 5).

**Figure 5.**
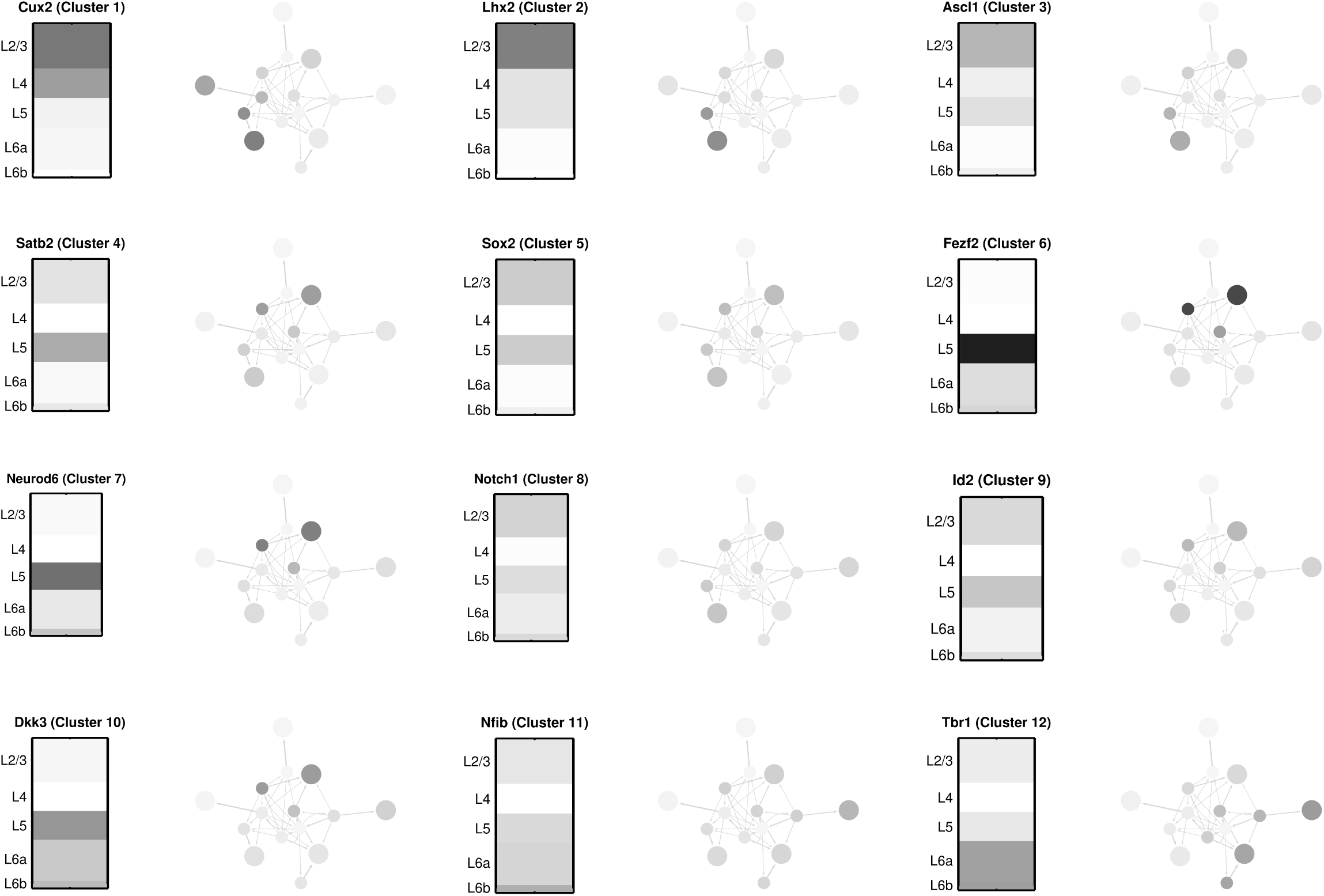
Prediction of transcription factor expression across precursors. The expression patterns of 1751 transcription factors was measured in the adult mouse cortex by Belgard et al. (2011). We clustered these patterns into 12 groups according to similarity of their laminar distribution (see Table S1). The expression pattern of one representative factor from each group is shown in the 12 schematic cortical columns (grey value in proportion to observed expression). For each case, the adult expression pattern was assigned to the terminal states of the *D* = 10 State Diagram (Figure 3). These values were propagated backwards into the SD as explained in the text. Grey shades of precursors indicate their predicted expression of that transcription factor. Thus, the 12 SDs together predict the profiles of expression of the 12 factors (and their groups) across all the cell states of neurogenesis as encoded by the State Diagram.

### 4.6 Abstract Gene Regulatory Networks

The second, complementary model, is functional. The states and state transitions are implemented implicitly by a *genotypic* model (or Gene Regulatory Network, GRN) (Figure S1C). In this case the interactions between genes and transcription factors are explicitly modeled. The network is designed in such a way that the global developmental process arises from the local dynamics of genes in individual cells. This model is visualized as a graph (not a tree), in which the nodes represent genes, and the edges represent interactions between genes. Importantly, the genotypic model is mechanistic in that it not only expresses allowable states and state transitions, but also declares the causal mechanisms by which the states are implemented, and reached.

### 4.7 An abstract GRN for mouse cortical neurogenesis

We will describe in detail below how the State Diagram (SD) can be estimated from experimental data, and how a GRN can be constructed that expresses this SD (and therefore the observed experimental data). Briefly, we first show that a low dimensional SD, composed of a small set of states, is sufficient to explain the generation of the different morphological cell types of the neocortex. This phenotypic model is then matched to a corresponding genotypic model. Because this problem is ill-posed (multiple genotypic models are able to explain a single phenotypic model), we restrict the domain of solutions by seeking a biologically realistic model based on a GRN. In our implementation, division asymmetry leads to differential inheritance of transcription factors in the daughter cells. This process is used to drive changing rates of cell numbers and types produced.

The SD generative model derived above is an example of a *phenotypic model* that describes the observed experimental data by assigning to each cell a state, and probability of transitions between those states at the time of cell division. This is essentially a phenomenological description of the statistics of neurogenesis. However, the question of the actual biological mechanism that expresses this statistical behavior is a much deeper one. Biological systems do not have a single constructor with global knowledge, able to direct all aspects of development. Instead, the only construction information available resides in the genetic instructions present in, and essentially localized to, each cell. The challenge then, is to implement the complex process of biological development as a *genotypic model* of neurogenesis. In this model developmental control is localized to gene regulation within individual cells (Figure S1C). The result of the operation of the GRN, distributed in its various configurations across all the lineages of neurogenesis, should be observable as the SD. Thus, we need to make the bridge from gene-level dynamics in individual cells, to the population-level stochastic behavior of the SD.

We have previously reported a formal language able to describe cellular and molecular processes that support cortical development (Zubler and Douglas, 2009). In particular, that language is able to control the development of a simple laminated cortical column (Zubler et al., 2013). However, in that previous work the generation of different cell types required precise ad hoc tuning of a system of differential equations. By contrast, our goal here was to create a genetic network model based on observed cellular mechanisms that is robust to intrinsic noise, reliable in execution, and flexible in the range of cell types it can generate.

The cellular machinery is composed of several layers of regulation. At the outermost layer, functional proteins fulfill specialized tasks such as structural support, movement, and cell morphology. Deeper in the regulatory machinery, DNA-binding regulatory proteins (transcription factors), define the progression through different cell activity states by regulating the gene expression profile of each cell. Transcription factors influence one another’s expression over time by binding to specific gene regulatory regions. The overall combination of the core regulatory network composed of transcriptions factors as well as the functional genes responsible for the cell phenotype, is referred to as a *Gene Regulatory Net-work* (GRN). However, the description below focuses largely on the transcriptional aspect of the GRN.

The concentration of each gene *x*_*i*_ is computed as a function of the concentration of other genes **x** = *x*_1_, *x*_2_, …, *x*_*n*_ by the rate equation:

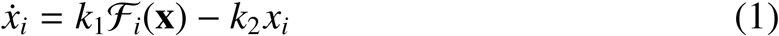

with:

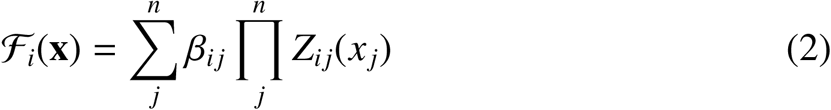

The function *ℱ*_*i*_(**x**), or *sigma-pi function*, is a linear combination of elements *Z*_*ij*_, each of which represents the binding of a transcription factor *j* on gene *i* as a function of its concentration *x*_*j*_ according to a sigmoidal probability binding function, the Hill function *Z*. Linear combinations of *Z* elements, determined by the coefficients *β*_*i j*_ ∈ {0, 1}, describe how transcription factors interact with each other by steric interactions. This formulation provides a model to express transcriptional networks as compositions of continuous Boolean logic gates (Figure S8), for which we propose an intuitive formal language based on logic gates.

Decisions leading to the acquisition of an appropriate cell fate rely on the ability of cells to commit to different stable states. A system that can perform such a task is a module with competitive and cooperative interactions. The most simple example of such a system is the bistable switch (Niwa et al., 2005; Huang et al., 2007), in which two auto-catalytic transcription factors *A* and *B* negatively regulate each others expression:

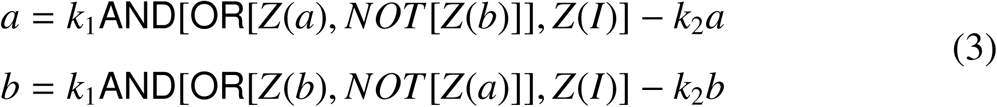

where *a* and *b* refer to the concentrations of the proteic product of genes *A* and *B*, and *k*_1_ = 1 and *k*_2_ = 1 represent production and degradation constants respectively. The system can be driven toward a specific state by an input *I* and is explicitly designed to display hysteric behavior upon input withdrawal: The network can remember the existence of past input signals (Figure S9). This design feature confers remarkable stability of the gene expression, and makes the dynamics of the module dependent only on an initial input signal (Jacob and Monod, 1961; Glass and Kauffman, 1973; Hartwell et al., 1999).

Biological development can be viewed as a sequential progression of precursors through different gene expression profiles; each cell state is associated with a characteristic profile. Thus, each lineage tree expresses one stochastic lineage of profiles arising from a given root precursor. The crucial question for understanding the dynamics of neurogenesis is how distinct profiles arise during the mitoses of the lineage, and so allow different fates for daughter cells. In our model this important property is due to possible differential distribution of transcription factors to the daughters. Each gene *X* is characterized by an asymmetry constant parameter *α*_*X*_, corresponding to the asymmetric division constant of its protein. Asymmetrical cell divisions lead to different distributions of transcription factors in the daughter cells, and thus to different gene expression profiles. Thus, cells regulated by a single bistable switch with asymmetry constants *α*_*A*_ and *α*_*B*_ can produce a range of cells with differing fates as a function of the division angle *ω*, the orientation of the mitotic spindle with respect to the internal distribution of substances (Figure 8). We set the required *α* for each substance in the bistable switch given a normalization constant *N*, such that −1 ≤ *α*_*X*_ ≤ 1:

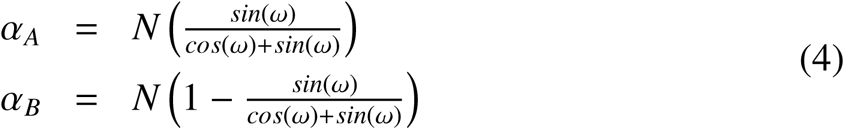

Beginning with the initial state “0” with low expression of both genes *A* and *B* (black cells), the activation of the input signal pushes cells to an undecided state “*AB*” characterized by high levels of *A* and *B* expression (orange cells). Either by the presence of an external influence, or by asymmetric cell division, cells can jump to states “*A*” or “*B*”, where only one gene of the bistable switch dominates the expression (pink or blue cells). Depending on the extent of the jump, each cell has a defined probability to reach new, otherwise inaccessible states. The irreversibility of jumps in the genetic landscape is implemented here as a dependency of the asymmetry constants on the gene product concentrations of the bistable genes. Once the motif reaches status “*A*” or “*B*”, further asymmetric division are inhibited, thereby limiting backward jumps to previous undifferentiated states.

The stochastic progression of precursors down differentiation paths can be modeled by a sequence of multiple genetic bistable switches, where each switch represents a branch in the differentiation decision tree and transition probabilities are mapped to cell division angle probabilities. Additional genes are required to detect specific transcription factor expression profiles and activate downstream functional programs. Control of precursor division is implemented by an independent clock mechanism that abstracts the complexities of the cell cycle and its phases. For simplicity it is assumed here to be a Gaussian distributed variable, independent on other events of the GRN. This basic genetic circuit is used to control cell fate decision at the moment of cell division, and to link the activation of different functional genes, such as genes responsible for cell migration, differentiation or apoptosis.

### 4.8 Self-construction of a volume of cortex in Cx3D

Finally, we validate the behavior of the GRN in a simulated physical environment using Cortex3D (Cx3D) (Zubler and Douglas, 2009), an agent and Java based simulation environment for investigating the physical growth of multicellular structures. This approach demonstrates the principles underlying the self-construction of a simple laminated cortical column and its neuronal connectivities (Zubler et al., 2013). In contrast to our earlier ad hoc system of differential equations for gene regulation (Zubler et al., 2013), we propose here a formal genetic language to design biologically plausible gene regulatory networks. We go on to demonstrate that the derived genetic network is able to control the generation of cortical laminae for different cortical areas by intrinsic genetic specification and by the information provided by the environment.

For the design of the GRN, sequences of bistable genetic motifs are used to encode cell fate decision at division and implement a genetic version of the state diagram for area 3 and 6. The SD was enhanced to introduce states for the generation of additional cell types (L1, subplate, and glial precursors cells), and to further reduce the overlap in the production of different cell types in time, as this has a dramatic effect on the stability of the simulation and the generation of homogenous layers.

Each state in the SD is mapped to 2 genes whose interactions implement the required bistable behavior. In addition, these genes are coupled to members of other bistable switches, or possibly to functional genes that execute cellular behaviors (Figure 6). State transition probabilities are encoded in the mitotic division angles that control the stochastic distribution of symmetric and asymmetric cell divisions. The core transcriptional network regulating the asymmetric distribution of cell fate determinants is composed of 36 genes. Further 24 housekeeping genes decode transcriptional expression into function, such as cell differentiation, migration, and other behavioral outcomes.

**Figure 6.**
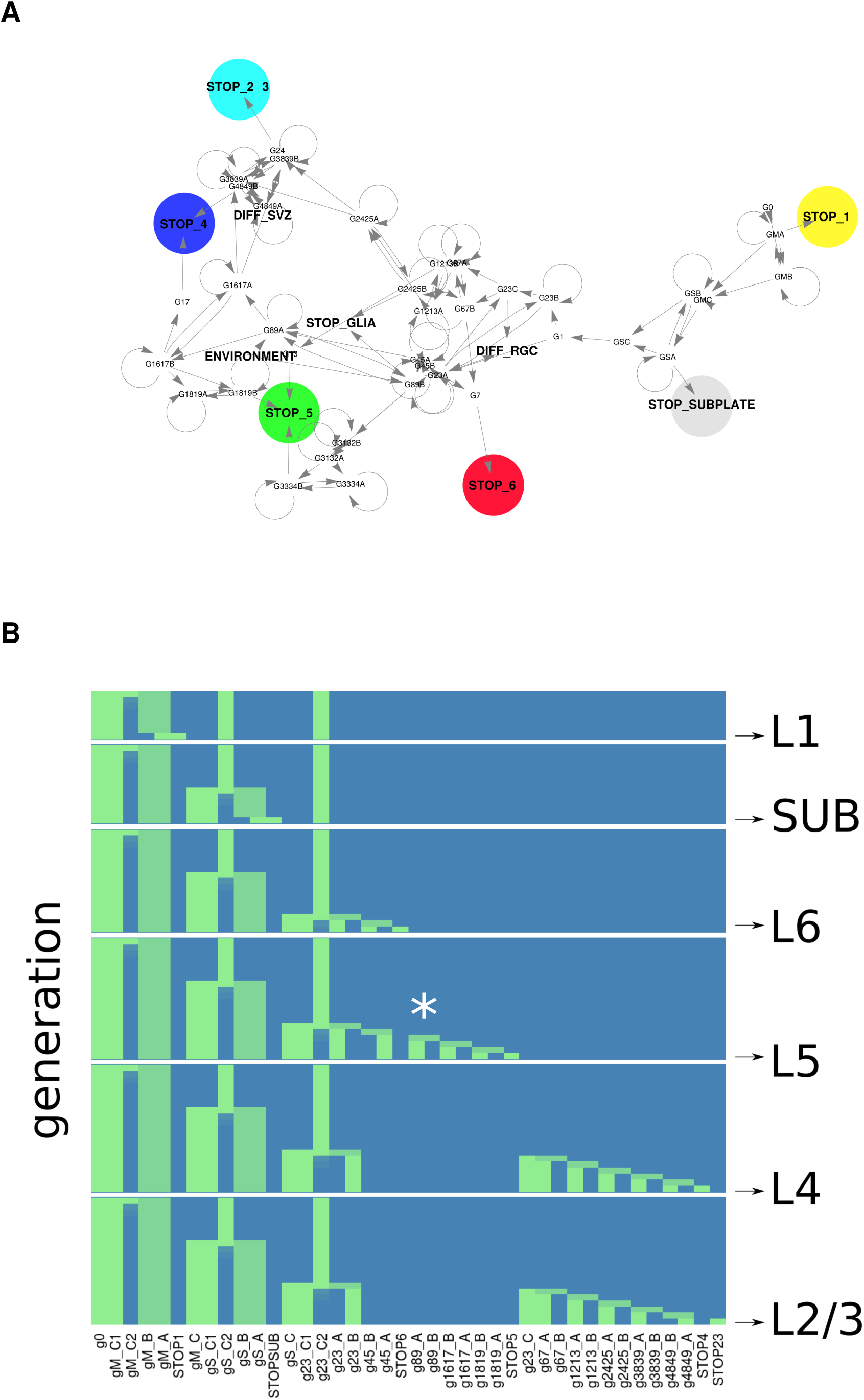
GRN controlling simulated development of mouse cortex. (**A**) Core Gene Regulatory Network controlling the production of marginal zone cells, and 5 different neuronal types of cortical area 3 and 6 in the mouse. Colored genes are expressed in neuron terminal states, and trigger differentiation. (**B**) Temporal expression pattern of core genes along lineages to 6 randomly selected cells of different type. Each panel shows the expression pattern of the initial precursor above, then patterns expressed by the next approximately 20 generations along lineage path, until terminal differentiating state is reached (below). Gene labels are shown beneath the lowest panel (L2/3). The expression patterns were measured immediately before mitosis, or at differentiation. At these times the genetic network reaches an attractor state. Expression levels range from 0 (blue) to 1 (green). Expression of gene ‘g89’, that biases neurogenesis towards either the area 3 or area 6 architectural phenotype, is indicated by white asterisk on path to layer 5 neuron.

The developmental model was then implemented in Cx3D (Figure 7). The simulation begins with an array of precursor cells in the neural epithelium lining the lateral ventricles (Figure 7, black cells). Each of these cell contains an identical copy of the genetic regulatory network (Figure 6A), initialized to its neuroepithelial precursor configuration. The precursors are aligned on the apical surface, and this orientation is used to establish the cell internal polarity axes.

**Figure 7.**
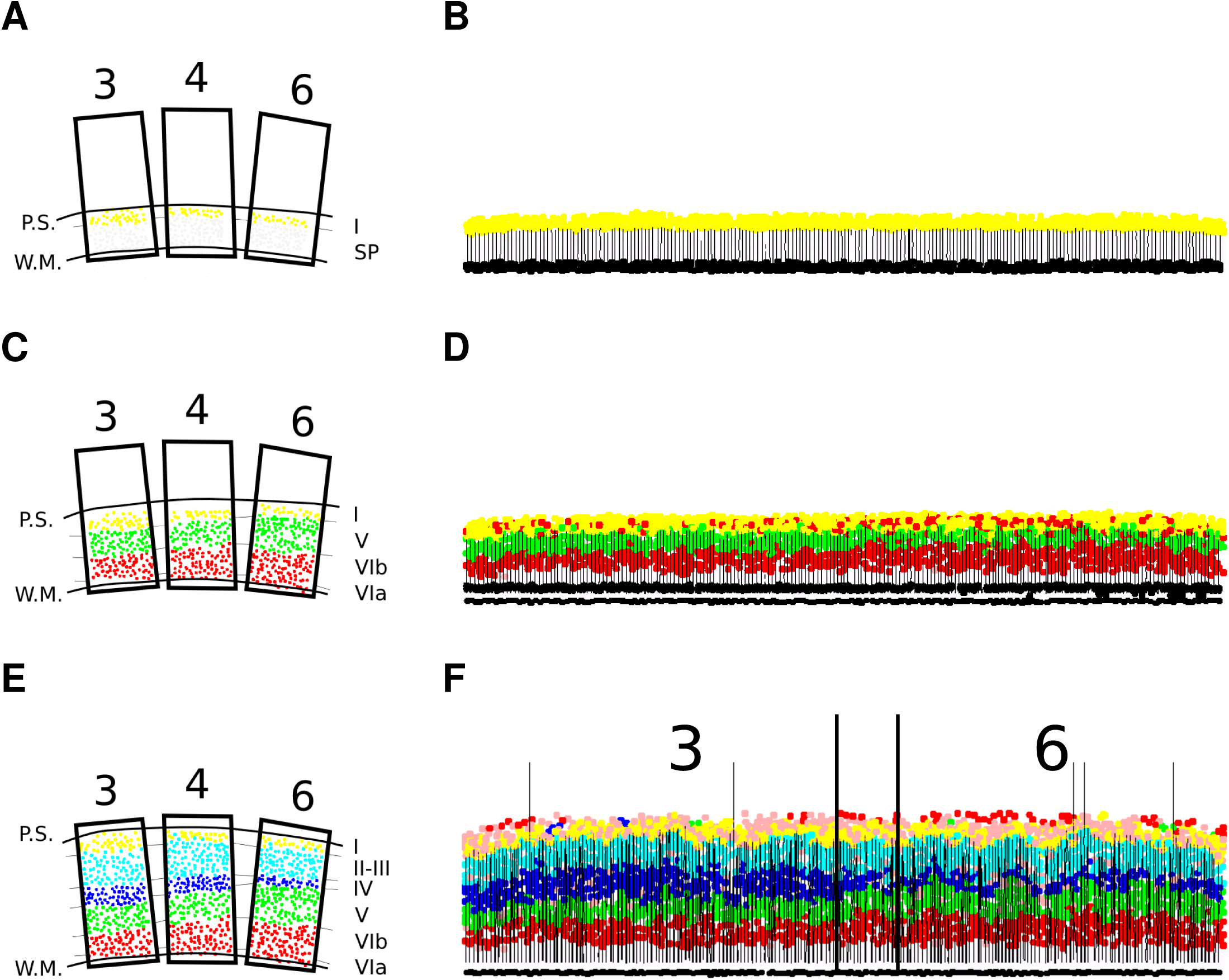
Simulation of cortical development. (**A**-**C**-**E**) Schematic visualization of cortical area 3, 4, and 6 derived from 500 *µm* paraffin sections counterstained with cresyl violet. Adapted from Polleux et al. (1997b). P.S., pial surface; W.M., white matter, SP, subplate. (**B,D,F**) Cx3D simulation of cortical development. For visualization, only a thin slice through the overall developing sheet is shown. (**B**) E11, with formation of marginal zone, subplate and radial glial cells; (**D**) E13, established infragranular layers; and (**F**) E16, established granular and supragranular layers, production of first glial cells. Area 3 and 6 boundaries marked by vertical black lines. There is a short transition zone between the 3 and 6 boundaries. Black: neuroepithelial cells; white/light gray: subplate cells; brown: intermediate precursors from subventricular zone; red: layer 6a and 6b; green: layer 5; blue: layer 4; cyan: layer 2/3; yellow: Marginal Zone or layer 1; pink: apoptotic cells; vertical lines, radial glia processes.

**Figure 8.**
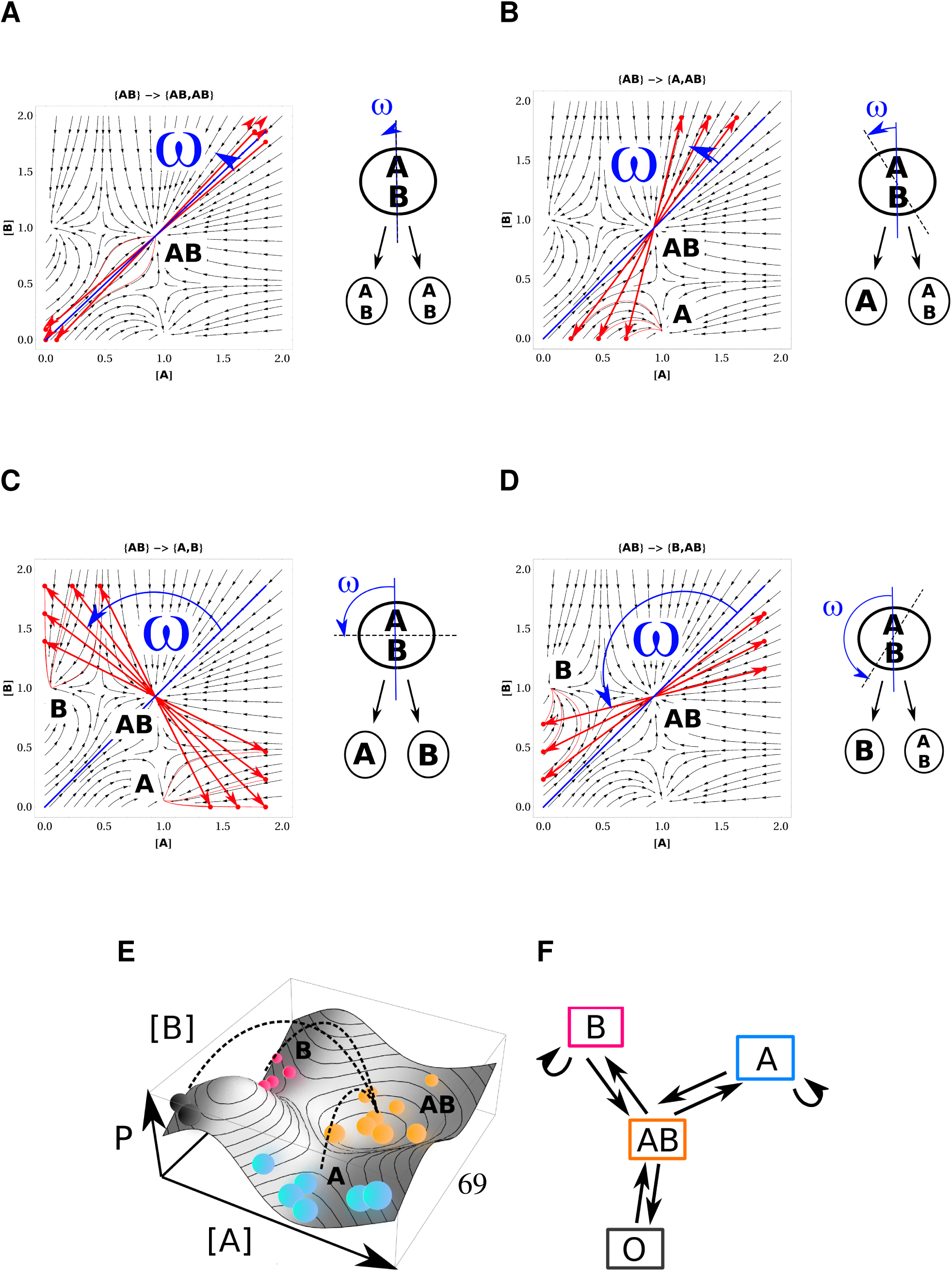
Genetic attractor landscape of a bistable switch with asymmetric cell division. Distributions of different division types as a function of division angle *ω*. Different division patterns arise: (**A**) {*AB*} → {*AB*}, {*AB*}; (**B**) {*AB*} → {*A*}, {*AB*}; (**C**) {*AB*} → {*A*}, {*B*}; (**D**) {*AB*} → {*A*}, {*AB*}. Red straight traces are simulated jumps at different angles, and red curvilinear trajectories show the time evolution after the jump. Blue lines indicate the *ω* angle with respect to the internal distribution of proteins. (**E**) Schematic representation of an attractor landscape *P* as a function of the concentrations of two genes *A* and *B*, in absence of an input stimulus. The landscape is determined by the manner of interaction between the genes. Each point on landscape corresponds to a possible gene expression profile. Spheres correspond to cells in different attractor basins; dotted lines to possible state transitions. (**F**) State diagram of bistable switch. Transitions are possible only by influence of the expression of another gene (e.g. through input *I*, Figure S9), or asymmetric cell division.

From this point onward, the behaviors of the distributed GRNs and the cells that they control are entirely autonomous. There is no intervention by a global controller, no explicit or global clock, and no explicit spatial coordinate frame. The only spatial cues are a pair of complementary morphogenic gradients in the medial/lateral axis of the neuroepithelial plate (Greig et al., 2013). The expression states of the distributed GRNs trigger their cells to undergo symmetrical or asymmetrical divisions according to their division angle, thereby forming the desired populations of successive precursors. The expression profiles at mitosis steer the stochastic transitions to successor states in the daughter cells. Mitosis is controlled by individual local cell cycle machines that induce cell cycle progression in precursors cells until they reach terminal differentiation. The entire process of neurogenesis from neuroepithelial cell to differentiated neurons involves some 20 mitotic divisions (Figure 6B).

Initially (E9-E12), the precursors progress through a sequence of increasing asymmetric divisions that lead to the production of the marginal zone (L1) and subplate cells, forming the early preplate. At the same time the VZ is formed. It is composed of radial glial cells (RGC) characterized by the extension of a radial process that often reaches the pial surface. Differentiating precursor cells that exit the cell cycle migrate along radial glial processes, constituting the successive waves of cell types that form the cortical plate in a inside-out manner. Migration is directed by local integration of guidance cues secreted by the marginal zone. A membrane bound stopping signal prevents cells from migrating past the pia. The density of cells in the marginal zone was also increased to provide physical containment of upwardly migrating cells.

In a subsequent phase (E13-E16) a second germinal layer, the SVZ is formed. In contrast to the VZ, precursor cells of this zone, the BPs, loose their radial process and apical polarity. In our simulation, lost processes are not degraded and continue to provide a scaffold along which neurons can migrate, increasing significantly the stability of the formation of distinct laminae. In this second phase, granular (L4) and supragranular (L2/3) are produced. The construction process ends with the establishment of the cortical sheet, and a residual germinal layer composed of glial cell precursors. Subsequently, corticogenesis would continue with a sequence of symmetric division for the generation of glial cells, and the growth of the first neural connectivities. These aspects are beyond the scope of the present paper, which is concerned only with the general principles of the GRN and its derivation.

The simulation exhibits a clear arealization of laminar organization that conform to the characteristics of areas 3 and 6 (Figure 7). The percentages of various neuronal types produced by the simulation in both areas also conform remarkably well to experimental observation (Table 1). There is a short intermediate zone between these two areas, corresponding to a cytoarchitectural boarder. This transition zone in the simulation may be analogous to area 4 that is interposed between areas 3 and 6 in mouse cortex, but which was not explicitly modeled.

**Table 1.** Laminar distributions of differentiated cells. Cells produced by simulations of GRN guided neurogenesis in areas 3 and 6. Quantification of simulated final neuronal production in each layer (before apoptosis) are compared with experimental data (Polleux et al., 1997a). Values are given in % with standard deviation. Experimental values were averaged and normalized to 100%.

| Layer | Area 6 |  | Area 3 |  |
| --- | --- | --- | --- | --- |
|  | Experimental | Cx3D | Experimental | Cx3D |
| **1** | $0.9 \pm 0.9$ | $13.7 \pm 0.0$ | $1.2 \pm 0.2$ | $13.7 \pm 0.0$ |
| **3-2** | $27.1 \pm 6.4$ | $23.8 \pm 3.8$ | $28.4 \pm 4.2$ | $22.1 \pm 3.5$ |
| **4** | $12.0 \pm 2.1$ | $12.3 \pm 2.7$ | $19.7 \pm 5.6$ | $20.5 \pm 3.1$ |
| **5** | $27.0 \pm 6.0$ | $24.9 \pm 3.1$ | $18.6 \pm 1.4$ | $17.8 \pm 2.3$ |
| **6** | $32.9 \pm 3.8$ | $27.0 \pm 4.1$ | $33.5 \pm 0.7$ | $26.0 \pm 4.5$ |

In the simulation, areal specificity is cued by the initial gradient of morphogens aligned with the medial/lateral axes of the developing sheet. The concentrations of these morphogens are transcription factors for a gene pair (‘g89A’ and ‘g89B’, Figure 6). These genes bias neurogenesis toward either an area 3 or an area 6 phenotype by slightly changing the distribution of the precursor pool, when threshold conditions on the morphogen concentrations are satisfied. The ‘g89’ is expressed on lineages leading towards L5 pyramidal cells. The onset occurs some 4 divisions before final differentiation, and there affects the relative generation of precursors fated towards layers 4/5. Thus, development towards area 3 or 6 occurs through a small and bias in the distribution of precursor cells, localized to particular region of the lineage tree (and so a time window) well before differentiation (Figures S5, 6B).

## 5. Discussion

We use ‘self-construction’ to refer to the process whereby a system is able to make use of physically encoded rules to steer its own elaboration, without the intervention of any kind of external supervisor. By contrast, ‘development’ refers to the biological process whereby a single, or small number of precursors replicate and differentiate toward a very large, diverse population of differentiated and functionally organized cell types. Thus, questions of self-construction are concerned with the abstract principles that underlie development of biological systems, but might equally well be applied to a future technology.

We choose to study biological self-construction in the neocortex, because cortical development presents many interesting challenges. For example, cortical neurons are produced far from their final location in the adult and so must undergo a long migration before they can complete their differentiation and formation complex long-distance connections. Further, the cortical construction process results in a rather uniform laminar sheet on which is superimposed a more detailed structural and functional arealization, suggesting that subtle modifications of a general process of neurogenesis may be sufficient to explain the apparent complexity of cortical neural circuits.

Cortical cytoarchitecture and its parcellation into distinct areas reflects the spatiotemporal modulation of neurogenesis (Dehay et al., 1993; Polleux et al., 1997a; Dehay and Kennedy, 2007; Rakic, 2009). From its simple origins as a single layer of proliferative cells in the embryonic dorsal ectoderm, the cortex grows through self-replication of a small population of precursor cells. The interplay between these many local mechanisms of cellular interaction, and their relationship to global system behavior, are easier to grasp through detailed models and their simulations (Fisher and Henzinger, 2007).

Here we have used a modeling approach to address the question of how a single cellular regulatory system could determine the generation of a diversity of neurons, including their laminar location. Of course, sufficiently detailed data describing the full mechanism of gene regulation and its consequences for the behavior of individual precursors underlying development are not yet available. However, we demonstrate here that it is possible to obtain substantial insight into developmental mechanisms using only sparse experimental data. With less than 40 genes we are able to recapitulate the steps of cortical development in silico with Cx3D.

Our approach has two phases. In the first phase the experimental data describing the generation of various neuronal types is used to estimate the stochastic SD governing the generation of possible cell lineage trees (*phenotypic model*). Then, in the second phase we implement the SD with a compact GRN-like state model (*genotypic model*) whose behavior then satisfies the experimentally observed dynamics of neurogenesis with quantitatively very similar cell distributions. This GRN is composed of abstract genes, whose patterns of expression determine the observed range of cell behavior.

### 5.1 State model of cortical neurogenesis

Hidden Markov Trees, which model Markov Tree processes over a set of trees of observed variables, and their conditional dependencies, have been used successfully to cluster cells and infer cell states from partial lineage tree reconstructions (Olariu et al., 2009; Pfeiffer et al., 2016). However, such inference requires a relatively large amount of data and is impractical for very sparse samples unless there are additional constraints on the probability distributions. Instead, we derived a lower dimensional representation of lineages using a simpler approach based on spectral clustering on graphs, whereby it is possible to exploit lineage information to cluster cells according to their phenotype, and that of their daughters.

We have introduced the concept of a SD to capture the complexity of the cell lineages. The SD model assumes that the underlying biological mechanisms can be modeled as a Markov process, according to which each cell, with its characteristic features, can be completely described by an unobserved state. The evolution of cell states is defined by the cell’s current state, which comprises the cell’s internal state and its immediate surroundings. In contrast to our related work (Pfeiffer et al., 2016) in which phenomenological data is used to classify progenitors cells in the primate cortex, we address here the use of genetic markers (transcription factors) to infer the probable developmental pathways followed by precursor cells until their terminal differentiation during murine corticogenesis.

Because we have only sparse data (i.e. we observe gene expression profiles on terminal cells only), we have used a simple approach based on spectral clustering, by which we cluster potential cell states according to the distributions of cell types that they are able to generate. The method was applied on cortical lineages inferred from experimental developmental data for areas 3 and 6. By this method we obtained a low dimensional age-dependent model that explains neurogenesis in both cortical areas, and which, in contrast to homogeneous Markov processes is able to explain this developmental process using only a restricted number of states and parameters.

The SD model predicts that already at the neuroepithelial stage the precursor pool may be somewhat heterogeneous in terms of their fate potential. For example multipotent progenitor cells may coexist with a more specific population of cell fate restricted cells, as suggested experimentally (Franco et al., 2012; Guo et al., 2013). Interestingly, because transitions in our model are stochastic, progenitors may exhibit some plasticity, including the limited ability to revert to less differentiated states. Such transitions have been observed recently in primate corticogenesis, but have not yet been observed in the rodent cortex (Betizeau et al., 2013).

Surprisingly, the models for adjacent areas display many similarities and few significant differences. Key parameters in a single GRN distinguish the specification of cortical areas 3 versus 6. This observation suggests the presence of *genetic control points*, that is a small set of genes whose expression is able to control the switch between alternative cortical developmental programs. This finding agrees with the observed molecular similarity reported in neighbouring areas of the human frontal cortex (Johnson et al., 2009). More generally, this property suggests that the many areas of cortex within a species, could be affected by the settings of a small number of parameters in an otherwise rather generic control structure in accordance with biological observations (Ng et al., 2009; Bernard et al., 2012; Hawrylycz et al., 2012). This discovery poses the questions whether the emergence in the evolution of the primate neocortex is also due to changes in few, key genes, which lead to the generation of a much complex and diversified cerebral cortex, and the significance of control points in biological processes in general (Dehay et al., 2015; Florio et al., 2015, 2016; Fiddes et al., 2018; Mitchell and Silver, 2018; Suzuki et al., 2018).

Obviously, the quality of the model depends strongly on the initial experimental classification of differentiated cell types, and a more extensive collection of data are required for a more precise version. In order to establish the general concept presented in this paper, we have relied heavily on the published cell birthdating data following pulse ^3^ *H*-thymidine injections made throughout murine corticogenesis (Polleux et al., 1997a). However the same principles can be readily applied to gene expression (e.g. Figure 5) and other phenotypic data (e.g. (Pfeiffer et al., 2016)) in future. While the recording in parallel of cell lineages and associated genetic markers is still a challenging technical endeavour, single cell tracking (Amat and Keller, 2013; Beattie and Hippenmeyer, 2017) or single cell profiling technologies (Bendall et al., 2014) would provide data at the necessary level of resolution.

### 5.2 Gene regulation by asymmetrical division

Our stochastic model of neurogenesis requires a number of distinct cell states in order to satisfy at least the experimental observations on which the model is based. The method of estimation of these states is constrained by additional more general structural knowledge such as the existence of lineage trees, binary mitosis, terminal states, etc. It is for this reason that it is possible to circumvent the seemingly ill-posed nature of moving from sparse data to an elaborate dynamical system that not only generates the original data, but will likely generalize to entirely different kinds of developmental data (e.g. gene expression, Figure 5).

The State Diagram alone provides a mathematical description of neurogenesis. However, it is difficult to relate that level of description to a biological mechanism. The most interesting and experimentally useful aspect of this paper is the recognition that it is possible to *implement* the global dynamics of a state model with plausible biological mechanisms that have implications for further experimental exploration. The implementation is based on basic cellular processes such as gene regulation, cell division, and asymmetrical repartition of cellular components. In particular, the importance of planar segregation of fate determinants during cortical developmental processes has been recognized experimentally (Noctor et al., 2008).

We employ the concept of genetic regulation using a gene network design based on small modules composed of bistable switches, each acting as an independent functional component. The importance of multi-stability and modular organization in molecular and genetic control has been recognized for over half a century (Delbrück, 1949; Jacob and Monod, 1961; Glass and Kauffman, 1973; Hartwell et al., 1999; Alon, 2006), however the modular networks reported here are arguably the largest such systems yet, that have been configured to control the development of complex tissue. We were surprised to find that the design of the GRN was less difficult than we had anticipated. Because the individual modules are functionally independent and self-restoring in their behavior, the interconnections between modules are rather insensitive to parameter settings. The overall network inside a given cell will converge toward its stable state, and it will finally trigger a mitotic division, though which it copies itself to its offspring. Thus reliable modules generate, by means of stochastic asymmetrical divisions, the desired distribution of cells over neuronal types. In this way, even an homogeneous pool of precursors can lead to the generation of diverse cell types. That is, the control of cell type and numbers is implicit in the asymmetric distribution of gene products, and how the genes influence one another’s expression.

Currently, the model GRN is composed of arbitrarily named abstract genes. Their significance rests only in that this set and their interactions are necessary to satisfy the expression states and transitions required to control the developmental process. The relationship between those model genes and actual experimentally named genes expressed in particular developmental systems needs to be comprehensively established. Establishing these relationships, as we have demonstrated by predicting the activation of transcription factors in the pool of precursor cells, and improving the model using the informative gene expression atlases will provide fruitful avenues for future research.

### 5.3 Simulation of cortical neurogenesis

The performance of the GRN was verified by simulation of neurogenesis using Cx3D (Zubler and Douglas, 2009). Cx3D respects physical processes such as mitosis, cell-cell interactions, movement and chemical diffusion in three-dimensional space. Each cell is an autonomous agent exerting only local actions, and using only locally available information. The physical behaviors of the cells are determined by the intracellular molecular processes expressed by the GRN. This large scale simulation of the physical mechanism makes it possible to bridge the scale between molecular processes and cell behavior.

The GRN is inserted into neuroepithelial prtecursor cells and initialized to a unique starting state. Each neuroepithelial cell contains also a simple cell clock that forces cells to divide at regular time intervals. Although the cell cycle length, in particular the length of the G1-phase, is correlated with the mode of cell division (Dehay and Kennedy, 2007; Pilaz et al., 2009; Lange et al., 2009; Arai et al., 2011) it was modeled here as an independent mechanism as the biological detail of this correlation is still unclear. The GRNs then orchestrate through their various stochastic expressions in the successively generated cells, different molecular and physical processes leading to cortical lamination. It is by virtue of asymmetrical division that progenitor cells undergo progressive cell fate restriction in accordance with experimental observations (Shen et al., 2006; Gaspard et al., 2008).

Modulation of only a single gene was sufficient to steer neurogenesis towards the characteristic architectures of either area 3 or 6. This finding suggests a generic developmental program for corticogenesis across the cortex, where a few localized factors elicit the differences in neuron number that characterize cortical areas. This locally modifiable generic program could account for the multiplicity of cortical areas, despite a relatively restricted number of transcription factor gradients in the early forebrain (O’Leary et al., 2007; Sur and Rubenstein, 2005; Greig et al., 2013). During evolution there is a progressive increase in the number of cortical areas reaching as many as 140 in macaque (Essen et al., 2011), despite an expected conservation of the early patterning of the forebrain (Donoghue and Rakic, 1999; Rash and Grove, 2006; Monuki and Walsh, 2001; Bayatti et al., 2008; Šestan et al., 2001; Sur and Rubenstein, 2005). It is likely that such a generic developmental program can be spatiotemporally modulated by extrinsic factors including afferent fibers originating from the sensory periphery as shown experimentally (Dehay et al., 1996; Dehay and Kennedy, 2009; Rakic et al., 2009; Krubitzer and Kaas, 2005), which coupled to genetic changes could lead to diverse evolutionary scenarios (Striedter, 2005).

We have shown in this paper that sparse phenotypic and cell lineage data can be used to derive an abstract GRN whose dynamics are able to control the detailed, quantitative, neurogenesis of the areas from which the original data was obtained.

The remarkable reliability of the modeled neurogenesis rests in the multi-stable and modular architecture of the GRN. Although mitosis may create offspring with different initial conditions, they will each reliably converge towards a permitted gene expression state and so to a recognizable precursor type of the cell lineage. Subtle and localized changes induced by mitosis in the stochastic distribution of transcription factors across offspring, can steer the overall profile of differentiated cells and their laminar location. The model can be used to explore and predict the forms of lineage and the resultant precursor pool sizes and relationships that precede the final adult cortical architecture.

While the present model of cortical neurogenesis is only an approximation to vast biological detail, is starts to explain the nature of the global coherence amongst multiple, distributed, locally independent cellular agents; and provides a useful tool for exploring the complex relationship between individual cell gene expression and population behavior underlying the development of the brain. Additionally it will also be a valuable tool for explaining diseases associated with gene regulation during cortical development.

## 6. Acknowledgments

We acknowledge helpful discussions with our SECO collaborators, in particular Kevan Martin, Christoph von der Malsburg, Michel Pfeiffer, and Adrian Whatley. This work was supported by European Union project grant FP7-216593 “SECO”. This work was also supported by LabEx CORTEX (ANR-11-LABX-0042)-HK, CD, and LABEX DEVweCAN (ANR-10-LABX-061)-CD of Université de Lyon, within the program “Investissements d’Avenir” (ANR-11-IDEX-0007) operated by the French National Research Agency (ANR); ANR-14-CE13-0036 (Primacor) and Fondation pour la Recherche Médicale (Equipe DEQ20160334943)-CD.

## 7. Methods

### 7.1 Cortical cell lineages reconstruction

We used published cell birthdate data from sensomotory cortex (Polleux et al., 1997a) to estimate the distribution of lineage trees underlying the neurogenesis of mouse area 3 and 6. Polleux et al. (1997a) employed pulse ^3^ *H*-thymidine injections made throughout corticogenesis to measure the variation of cell cycle duration, cell cycle exit probability *k*_*Q*_(*t*), and laminar fate *k*_*QX*_(*t*) as functions of developmental time *t*. Following their data and model we computed the temporal generation of neuronal types by numerical solution of the continuous differential equations describing cell proliferation and differentiation (Polleux et al., 1997b). We used these population distributions across developmental time to generate probabilistically instances of cortical cell lineage trees (Figure 1).

Cell proliferation can be seen as a discrete branching process whose time step Δ*t* is equal to the cell cycle length. At each time step, cells either differentiate terminally with probability *p*_1_ = *k*_*Q*_(*t*), or they divide with probability *p*_2_ = (1 − *k*_*Q*_(*t*)) to form two daughter cells. These possibilities can be represented formally by the probability-generating function (pgf) (Bremaud, 1988):

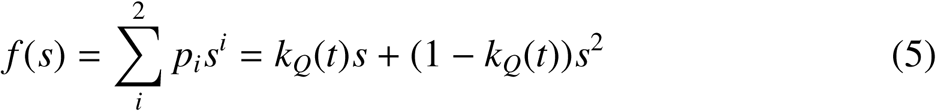

where *p*_*i*_ is the probability that a cell gives *i* offspring in the next generation and *s*^*i*^ is a dummy variable that accounts for the different numbers of cells generated. The pgf enumerates all the possible outcomes after one time step, and has the property Σ*_i_ p_i_* = 1. We used this formula recursively to generate possible sequences of cells from single precursor cells. Sixty probabilistic lineage trees were computed for each of the two areas.

### 7.2 Graphical representation of the State Diagram

The State Diagram (SD) describes the states of cells that appear in the CLT, and the genealogical relationship between these states. For each state there is a corresponding vector of observed features ⟨*f*_1_, *f*_2_, …, *f*_*L*_⟩. States for which features have been observed experimentally are defined as labeled, otherwise the states are unlabeled or hidden. We assumed that observed features (e.g. neuronal morphologies, gene expression) are available only for terminal cell states, and that the features of all the precursors are hidden.

It is convenient to represent the State Diagram in the form of a directed graph. Recall that *𝒢* = {*𝒱, ℰ*} is a directed graph with vertices *𝒱,* = {*v*_1_, *v*_2_, *…, v_n_*} and directed edges *ℰ* = {*e*_*i j*_} ⊆*𝒱* × *𝒱*. In a weighted graph, each edge is assigned a specific value, its weight. For such weighted directed graphs, there is an asymmetric, non-negative adjacency matrix **W** that associates each edge with a weight as following: *w*_*i j*_ = 1 if there is a direct link that connects node *i* to node *j* or *w*_*i j*_ = 0 otherwise. Also, we define the *in-degree* matrix *D*_*in*_ as the diagonal matrix of the sum of weights on incoming edges and the *out-degree* matrix *D*_*out*_ as the diagonal matrix of the sum of weights on outgoing edges:

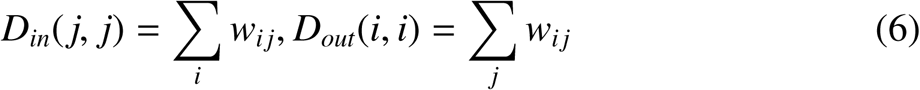

Given a directed weighted graph, there is a natural random walk on the graph defined by a transition probability matrix **P**, where *p*_*i j*_ = *w*_*i j*_/*d*_*out*_(*i*) for all edges, and 0 otherwise. Thus, in this naive random case, transitions on the outgoing edges are equally probable, and sum to 1. The situation for the State Diagram is somewhat different. Each vertex *V* of the State Diagram corresponds to a cell state, and each edge *E* asserts a genealogical relationship between connected states. Now the transition probability matrix *P* represents the strength of these genealogical paths between states. That is, it represents the proportion of cells in the source state that will undergo each of the allowable transitions, multiplied by 2 to account for the doubling of cell number by mitotic division. *P* must be estimated from data.

### 7.3 Dimensionality reduction of the State Diagram

Given an SD and vectors of observed features ⟨ *f*_1_, *f*_2_, …, *f*_*L*_ ⟩ for its labeled terminal nodes, we consider the task of computing a pairwise similarity measure between all nodes of the SD based on how unlabeled nodes are connected to labeled nodes. For undirected graphs, a widely used method for computing structural similarity is spectral clustering (Chung, 1997; von Luxburg, 2007). This method makes use of the spectrum (eigenvalues) of a similarity matrix to cluster data into groups of highly similar nodes. For our case of directed graphs, we introduce an approach based on the Laplacian **L** of the normalized directed matrix:

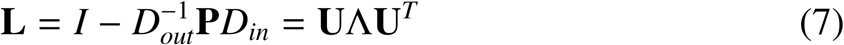

where *P* is the directed transition probability matrix, *D*_*out*_ is the out-degree matrix, *D*_*in*_ is the in-degree matrix, and *I* is the identity matrix. Λ = *diag*[*λ*_1_ ≤ *λ*_2_ ≤ … ≤ *λ*_*n*_] is the diagonal matrix of eigenvalues, and **U** = [**u**_1_**u**_2_ *…* **u***_n_*] is the orthonormal matrix with eigenvectors of **L** in each column. **U** : 𝒱 → ℝ*^n^* provides an embedding of each vertex in an *n*-dimensional metric space. Each column of **U** corresponds to an axis of the space, while each row of corresponds to the coordinates of a vertex in that space. The Euclidean distance *δ* between pairs of nodes (*r, s*) provides a distance matrix:

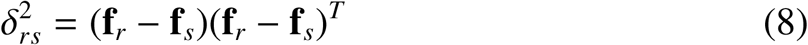

Mapping of the State Diagram to a *n*-dimensional space is particularly useful, because conventional algorithms such as hierarchical clustering can be applied there. We used the single linkage algorithm to perform clustering on the distance measure. Nodes whose distance was less than a specified threshold were clustered into a single node, which was assigned the average of their transition probabilities. The projection is in Euclidean space and so the feature vectors for each clustered node can be computed by solving a linear equation, because we assume that each node can be represented by a linear combination of feature vectors:

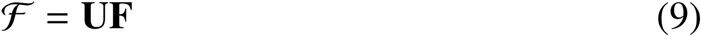

where **F** is a *n* × *l* matrix containing the features of the observed states, **U** is a *n* × *n* matrix, and *ℱ* is a *n* × *l* matrix with observed and estimated features. For visualization purposes, each terminal state was also matched to a 3-element feature vector **F***_RGB_* representing a unique color, and colors of all states were estimated by *ℱ*_*RGB*_ = **UF**_*RGB*_.

We validated our spectral clustering method by measuring its performance on a set of artificial lineages generated by ‘ground truth’ models. The classification of cells to states by the algorithm was compared against 100 deterministic, stochastic and random cell lineages each composed of 5 states. The fraction of states misclassified by the algorithm are shown in the confusion matrices of Figure S3. The columns of the matrices represent instances of predicted states, while the rows represent instances of ground truth states. We found that deterministic ground truth models are recovered in 100% of cases, while probabilistic ground truth models are recovered in 80%. This decrease in performance on probabilistic models is due to misclassification of states as well as to the existence of multiple equally likely solutions. The chance of random prediction of 5 states is estimated at 18%. These results demonstrate that a low dimensional SD can indeed capture the statistical variation of the cell lineage data at above chance level.

### 7.4 Multi-type Markov Branching Process

A State Diagram can be interpreted as a Markov branching process with multiple states. A branching process is a discrete-time random process that models a population in which each particle in generation *t* produces some number of individuals in generation *t* + 1, each of which can assume one of *m* different states.

Let *S* denote a finite set of states *S* = {*s*_1_, *s*_2_, *…, s_m_*}, and *Z*_*n*_ = (*z*_1_, *z*_2_, *…, z_m_*) the vector of variables describing the population size at the *n*’th generation in each state. Then the time-invariant transition probability *p*_*i j*_ describes the probability that a particle will transit from state *i* to state *j* (Markov property):

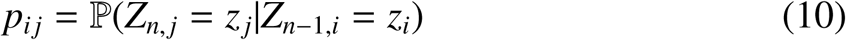

The system evolution is completely characterized by the set of states, the marginal distribution of its initial state *Z*_0_, and the transition probabilities between states. We write the joint probability distribution of *Z*_*n*_:

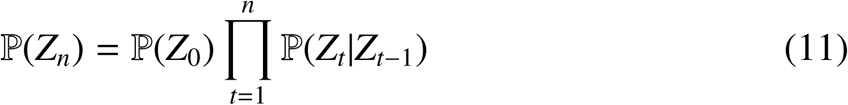

By setting the elements of the weight matrix *P* equal to the probability of moving from state *i* to a state *j*, the equation may be rewritten in matrix representation:

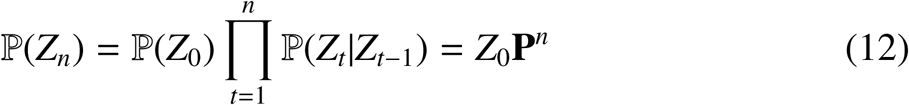

Markov models have limited ability to describe complex time-dependent processes using only a restricted set of states. Therefore, we extended this homogenous Markov model (HM, probability **P**) by two further approaches. First, as a non-homogeneous model (NM, age-dependent probability **P**(*a*)). Here each state transition probability is multiplied with an additional parameter that is set to 0 once a maximal number of self-replicating divisions is reached. This has the effect of truncating the long tails that are characteristic of Markovian processes. Second, as a time-dependent model (TM, time-dependent probability **P**(*t*)) that explicitly encodes the state transition probabilities for each time point. In order to compare branching processes for these three different approaches and different model dimensions, we computed their errors as the number of misclassified cells (cells in wrong terminal states) over the total number of cells produced at the end of the developmental process.

### 7.5 Formal genetic language definition

We designed a genetic “language” in order to describe gene regulatory networks (GRNs). This language was based on a set of variables *x* ∈ ℝ _≤0_ that represent substance concentrations, and a set of allowed operations on the substance con-centration values. This formalism greatly simplifies the construction of GRNs for developing systems as it is based on the design of the network topology, so that parameter tuning is reduced to a minimum. Although abstract, the formalism can be cast directly into the corresponding kinetic differential equations:

#### Read

Information about transcription factor concentrations is obtained from the environment through the Hill function *Z*, which computes the binding probability of a transcription factor to a promoter region given affinity constant *θ*, cooperativity *m* and binding bias *b*.

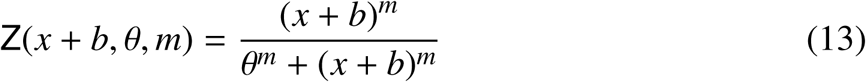

#### Write

Information can be written to the environment by the production of a given substance according to the rate equation, which influences the current substance concentration. *ℱ* takes the form of one of the possible logic operations, or combinations thereof.

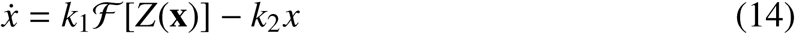

#### Distribute

Information is encapsulated by the cell membrane, which prevents external agents from directly interacting/modifying the cellular molecular components, and so provides a protected environment in which the cell performs its local computation. During development, a cell *c* divides and distributes its internal components asymmetrically to daughter cells 2*c* and 2*c* + 1.

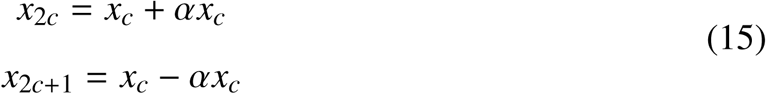

#### Logic operations

Logic operations are used to compute the result of the binding of multiple transcription factors to the promoter region, where *y*’s can be either the output of *Z* or the output of another logic operation.

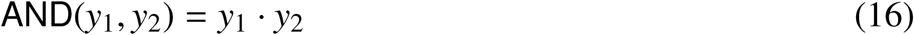

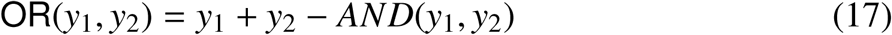

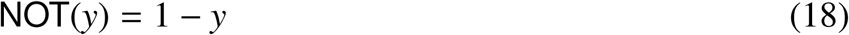

#### Derived logic operations

The elementary operations can be composed into derived operations, for example:

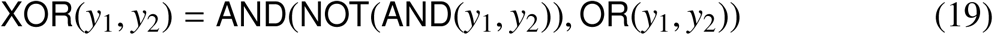

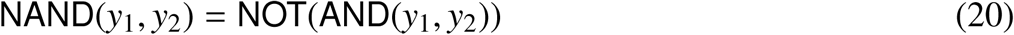

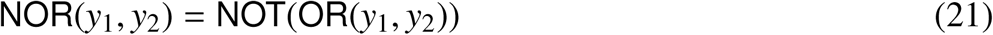

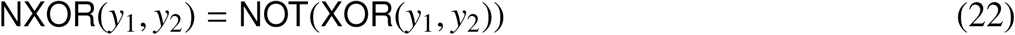

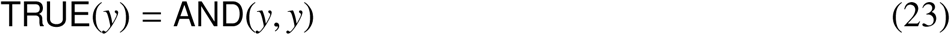

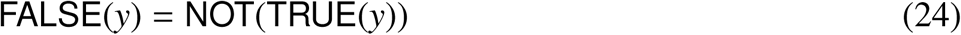

Another useful derived operation is the threshold function Z*_o_*, that indicates a threshold at any desired value *tr* ∈ [0, 1]:

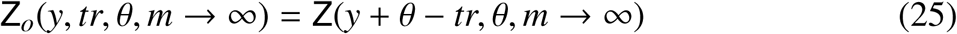

Notice that for co-operativity *m* → ∞, values of *x* are bounded to the set {0, 1}, logic operations behave as Boolean logic gates, and the genetic language reduces to conventional Boolean algebra.

### 7.6 Software

Spectral clustering was implemented in Matlab R2012a. Graph visualizations were performed using a Cytoscape 3.0 plugin (DynNetwork). Cortical simulations were performed using Cortex3D (Cx3D) (Zubler and Douglas, 2009).

## 9. Supporting Information: Tables

**Table S1.**
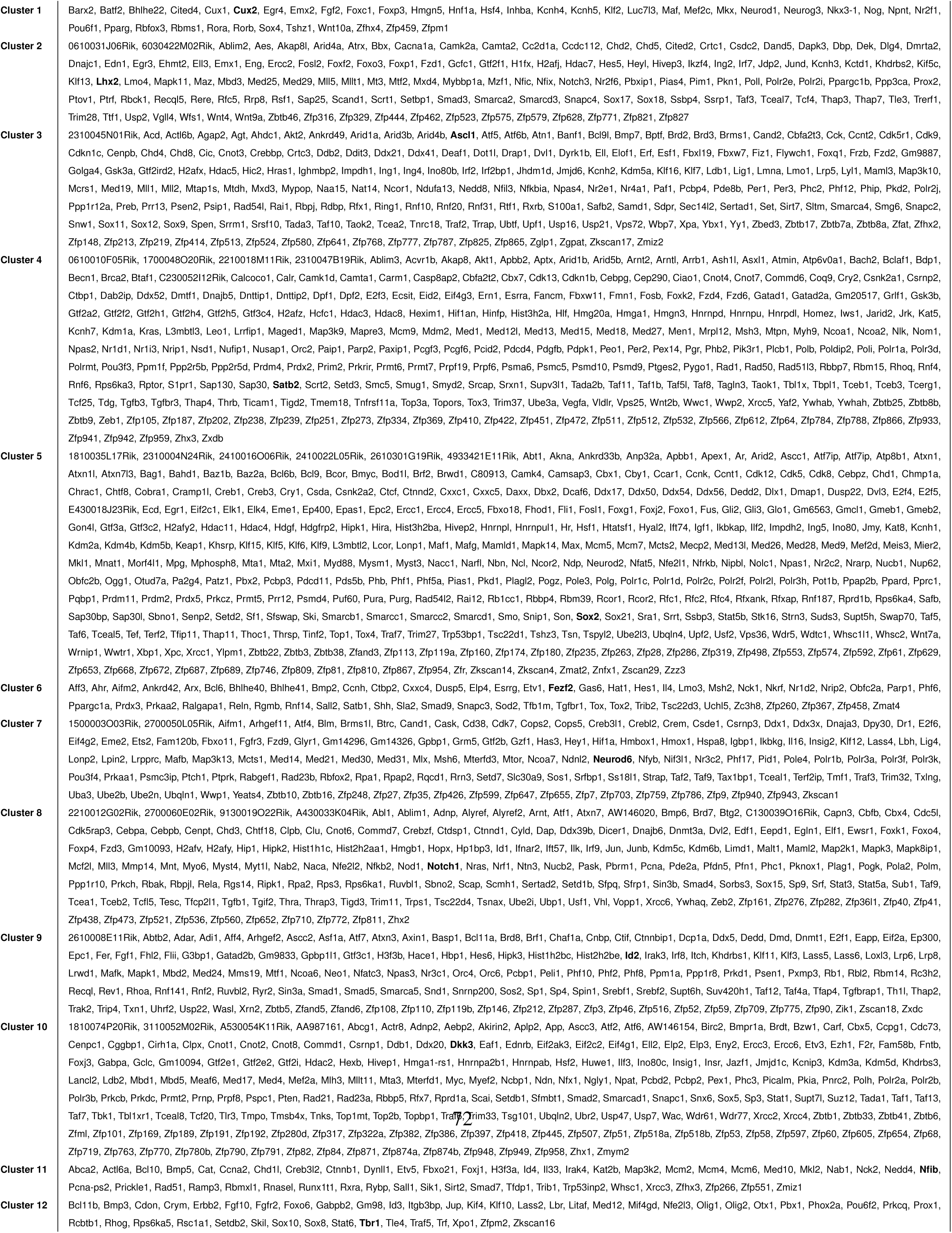
**Transcription factor clusters** 751 Transcription factors (Belgard et al., 2011) were clustered according to the distribution of their normalized expression values across layers 6a, 6b, 5, 4 and 2-3. The transcription factors of each cluster that were chosen as representative examples for Figure 5 are highlighted in bold.

### 10. Supporting Information: Figures

**Figure S1.**
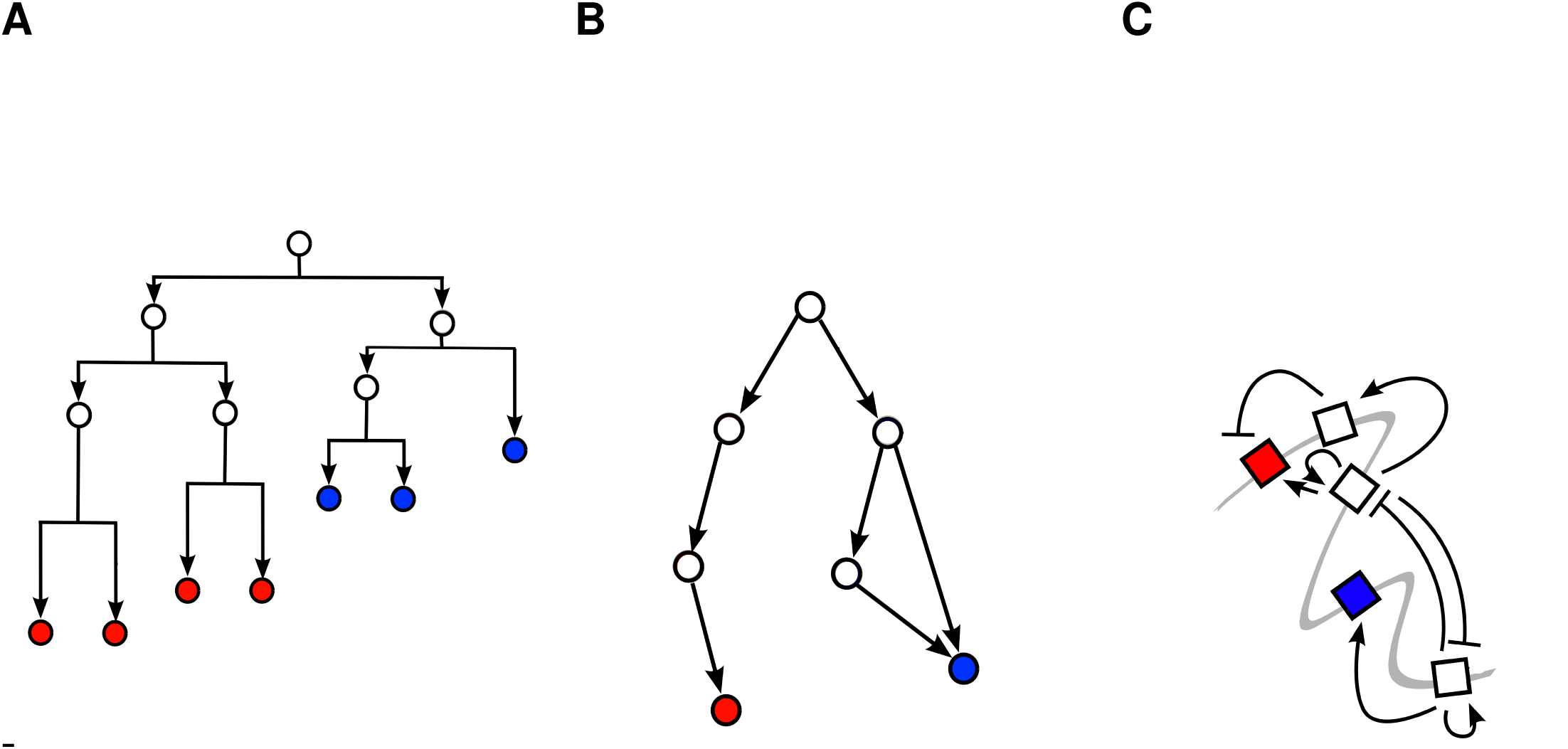
Aspects of biological development. The process of development can be understood in terms of three complementary models (**A**) The cell lineage tree describes the mitotic process rooted in a given precursor. Each cell divides symmetrically or asymmetrically to produce two similar or dissimilar daughter cells. Colors denote the different fates of terminal cells. (**B**) A phenotypic model of the possible states taken by cells of lineage tree. Each node represents a cell state that is characterized by a vector of observable features. Each edge represents a possible transition route between states. Colors denote the features expressed by terminal cell. (**C**) A genotypic model that is the mechanism underlying the lineage tree description, or the state diagram description. Each cell state is encoded by the expression of a subset of genes (squares) layed out on the DNA (gray line). The progression through the successive cell states of the lineage tree is controlled by gene interactions (black lines), and the degree of asymmetrical of cell division and gene interactions (black lines). These interactions may be positive (arrow) or negative (plate) with respect to their target genes. Colors represents genes linked with a particular terminal cell type.

**Figure S2.**
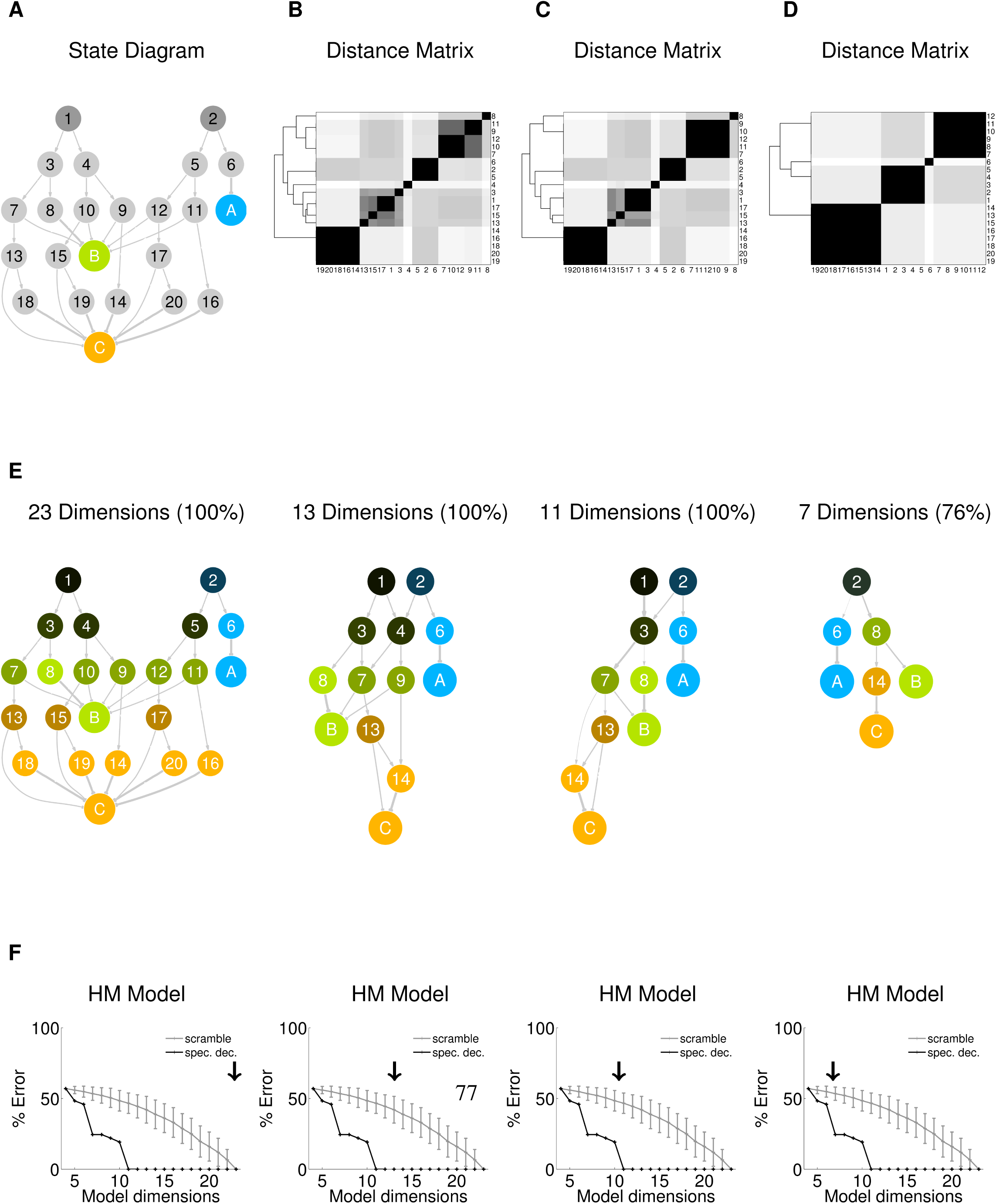
Reduction of State Diagram to lower dimensionality. (**A**) State diagram of example lineages (as Figure 2B). Nodes represent cell states, arrows state transition probabilities. States are labeled according to 3 observed features: *A* = ⟨1,0,0 ⟩ (blue), *B* = ⟨0,1,0 ⟩ (green), *C* = ⟨0,0,1 ⟩ (orange), and # = ⟨?,?,? ⟩ (gray) for states with hidden features. Initial states are depicted in dark gray. (**B**-**D**) State clustergrams of computed distance between every state pair with dimensions *D* = 23, *D* = 13, *D* = 11, and *D* = 7 (percentage of data represented in parenthesis). Dendrograms indicate hierarchical binary linkage of states. (**E**) Spectral label propagation on models, where each hidden node is colored according to its estimated feature distribution. (**F**) Model error as percentage of the correct final cell state distribution for spectral clustering (black) versus random model (gray, standard deviations for 100 trials). HM, Homogeneous Markov model.

**Figure S3.**
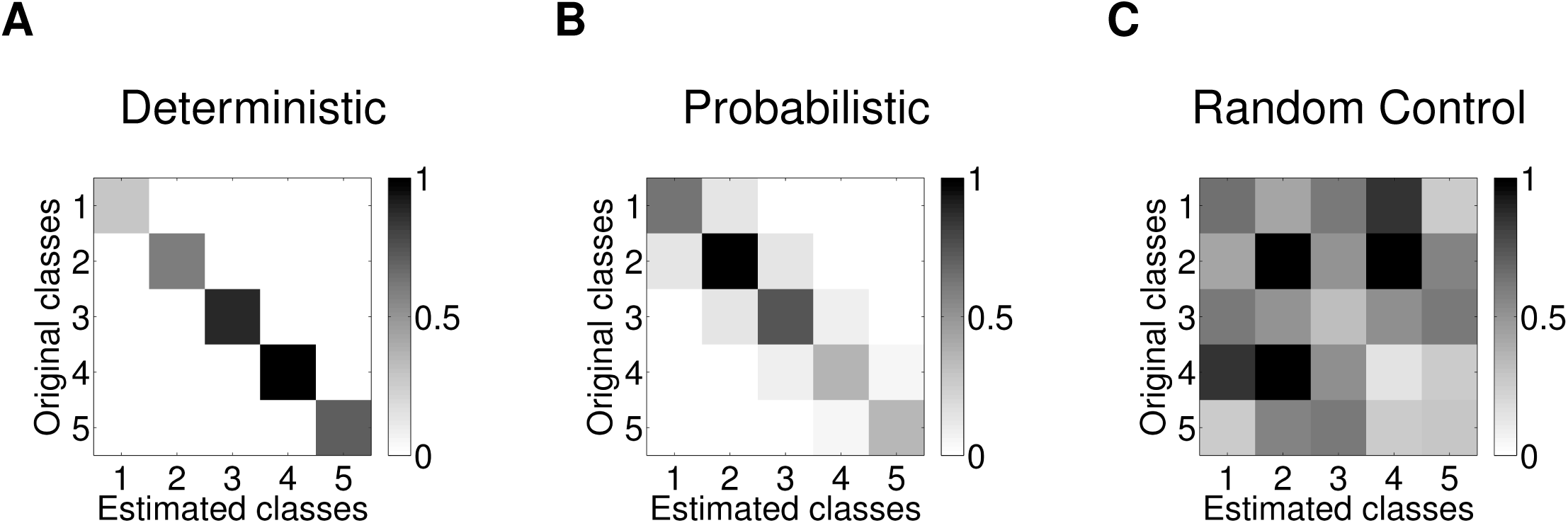
Classification performance of spectral clustering. The ability of spectral clustering to recover the correct Markov branching process was assessed on 100 lineages generated with 10 random 5-state models. Spectral clustering assigns a unique class to each cell, which is then compared to the known model class. (**A**) Confusion matrix of spectral clustering on deterministic model (0 ± 0% classification error). (**B**) Confusion matrix of spectral clustering on probabilistic model (20.3 ± 17.8% classification error). (**C**) Confusion matrix of random model (88.2 ± 18.7% classification error).

**Figure S4.**
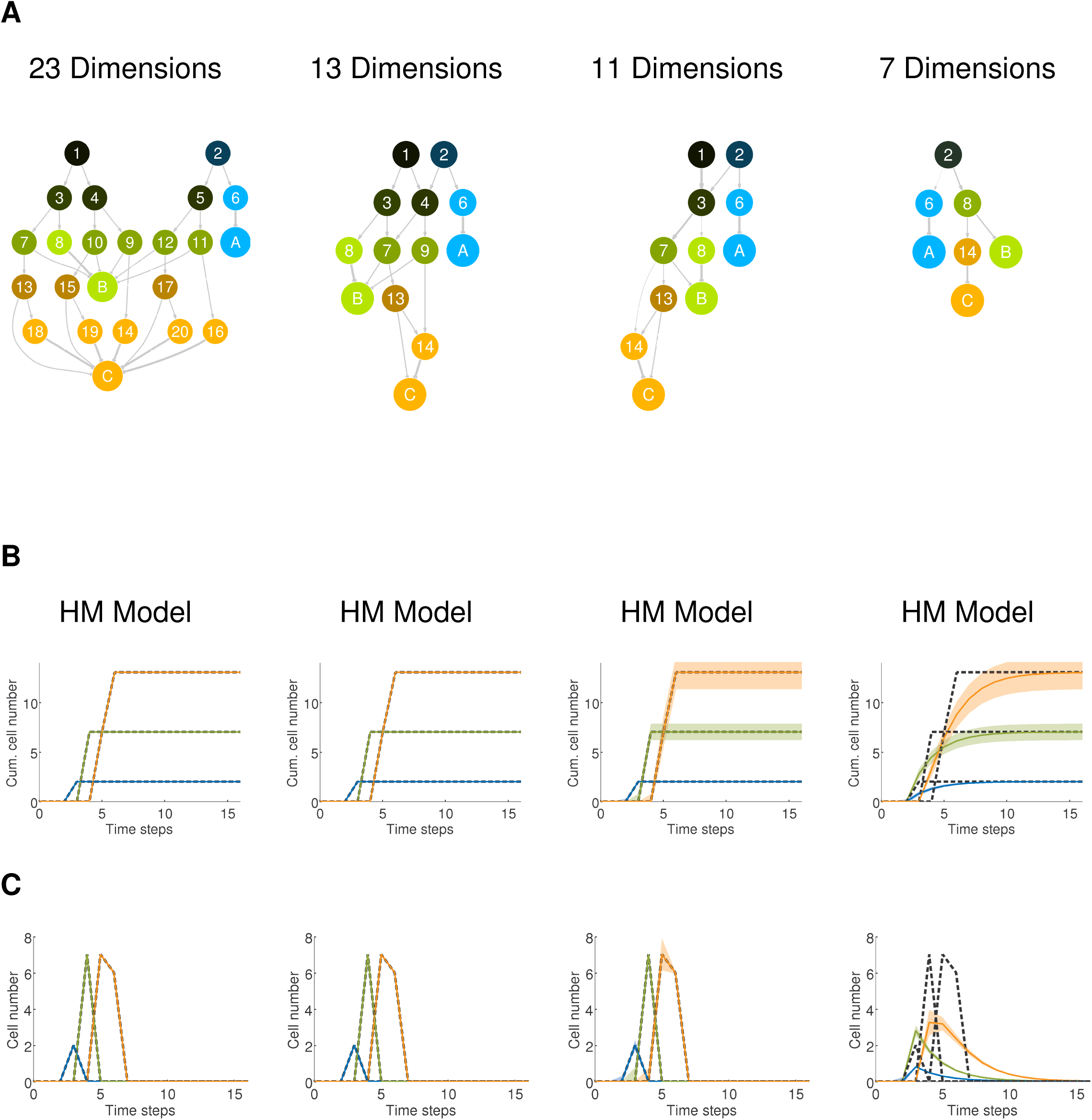
Cell type distributions generated by a State Diagram of decreasing dimensionality. (**A**) A State Diagram of an example sublineage is progressively reduced from dimension *D* = 23 to *D* = 13, *D* = 11, and *D* = 7. Nodes represent cell states, arrows state transition probabilities. (**B**) Output generated by Hidden Markov implementation of a State Diagram. Mean cumulative number of differentiated cells produced at each time step. (**C**) Mean instantaneous number of differentiated cells produced at each time step. Dashed lines, original distribution; colored lines, model distribution; shaded area, standard deviation. The *D* = 7 model fails to capture the original data.

**Figure S5.**
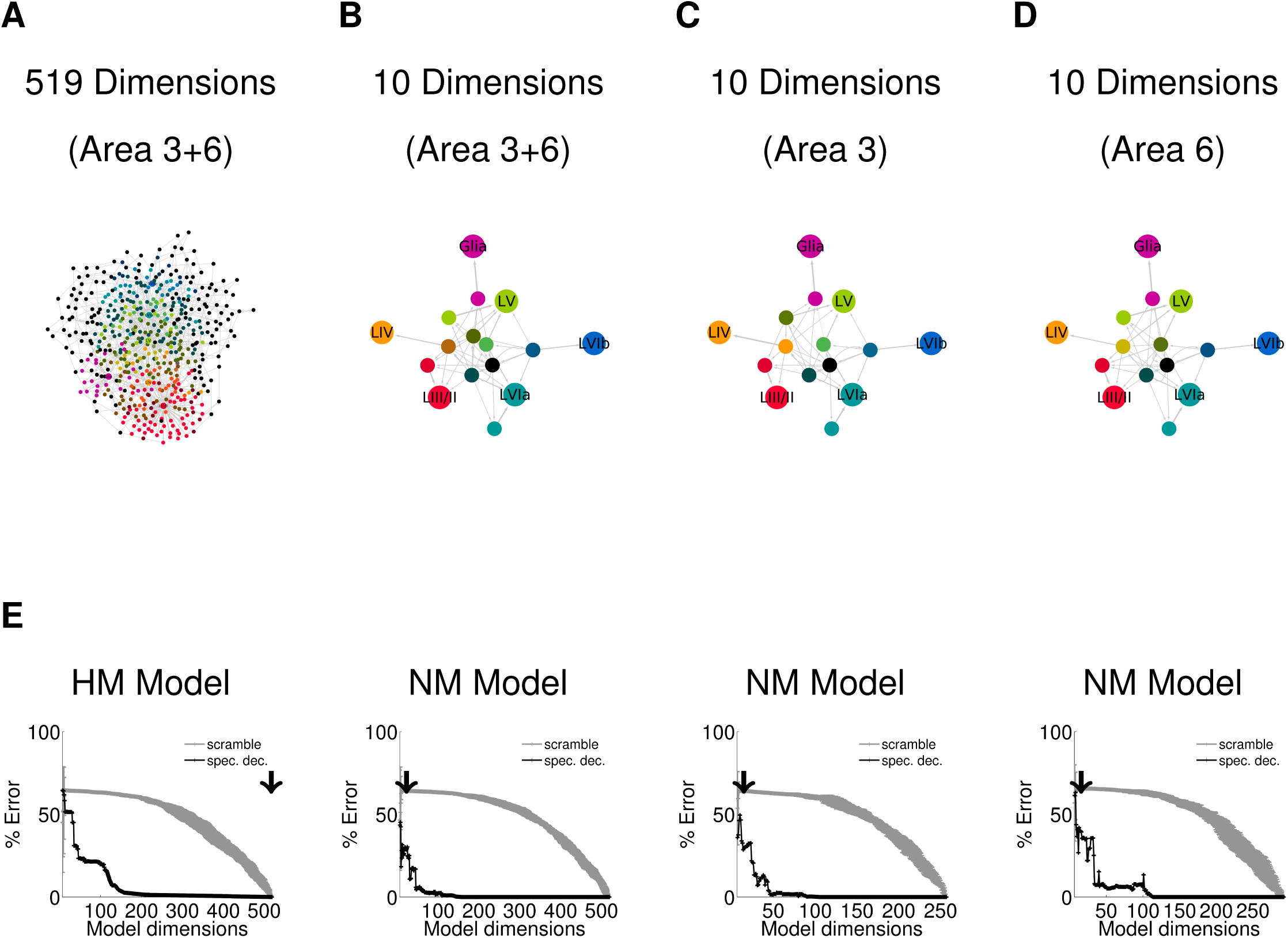
State Diagrams areas 3 and 6 combined, and separated. (**A**) 519-dimensional State Diagram of combined lineages for area 3 and 6. Nodes represent cell states, arrows state transition probabilities. (**B**) Combined SD reduced from *D* = 519 to *D* = 10 (area 3 and 6). (**C**) *D* = 10 SD for area 3 alone. (**D**) *D* = 10 SD for area 6 alone. Cell states: Layer 6b, blue; Layer 6a, sea green; Layer 5, green; Layer 4, orange; Layer 2/3, red; Glia, pink; and Unknown, gray. (**E**) Performance (% error against original data) of stochastic generator models (black traces) corresponding to the SDs above. The performance of the stochastic models is compared against a model free random control (grey traces). HM, Homogeneous Markov model; NM, Non-Homogenous Markov Model. Model dimension indicated by black arrow.

**Figure S6.**
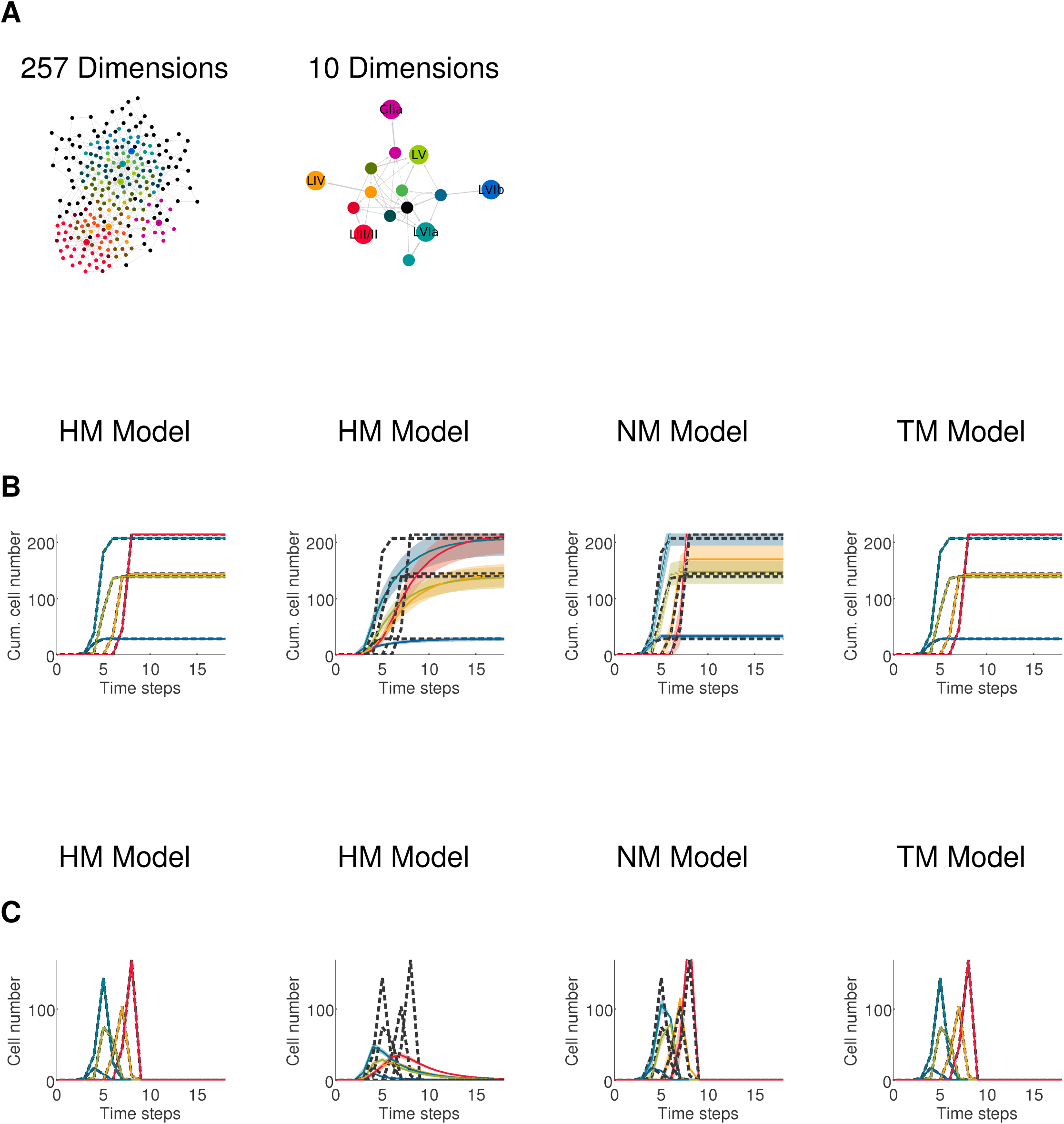
State Diagrams and model generated cell distributions for cortical area 3. (**A**) Original State Diagram *D* = 257 and its reduced *D* = 10 version for cell lineages in cortical area 3. Nodes represent cell states, arrows state transition probabilities. Cell state colors are the same as for Figure S5. (**B**) Generation of cells by various stochastic models. Mean cumulative number of differentiated cells produced at each time step. (**C**) Mean instantaneous number of differentiated cells produced at each time step. Dashed lines, original distribution; colored lines, model distribution; shaded area, standard deviation. HM, Homogeneous Markov model; NM, Non-homogeneous Markov model; TM, Time-dependent Markov model. Low-dimensional HM model fails to capture the data, whereas TM performs well.

**Figure S7.**
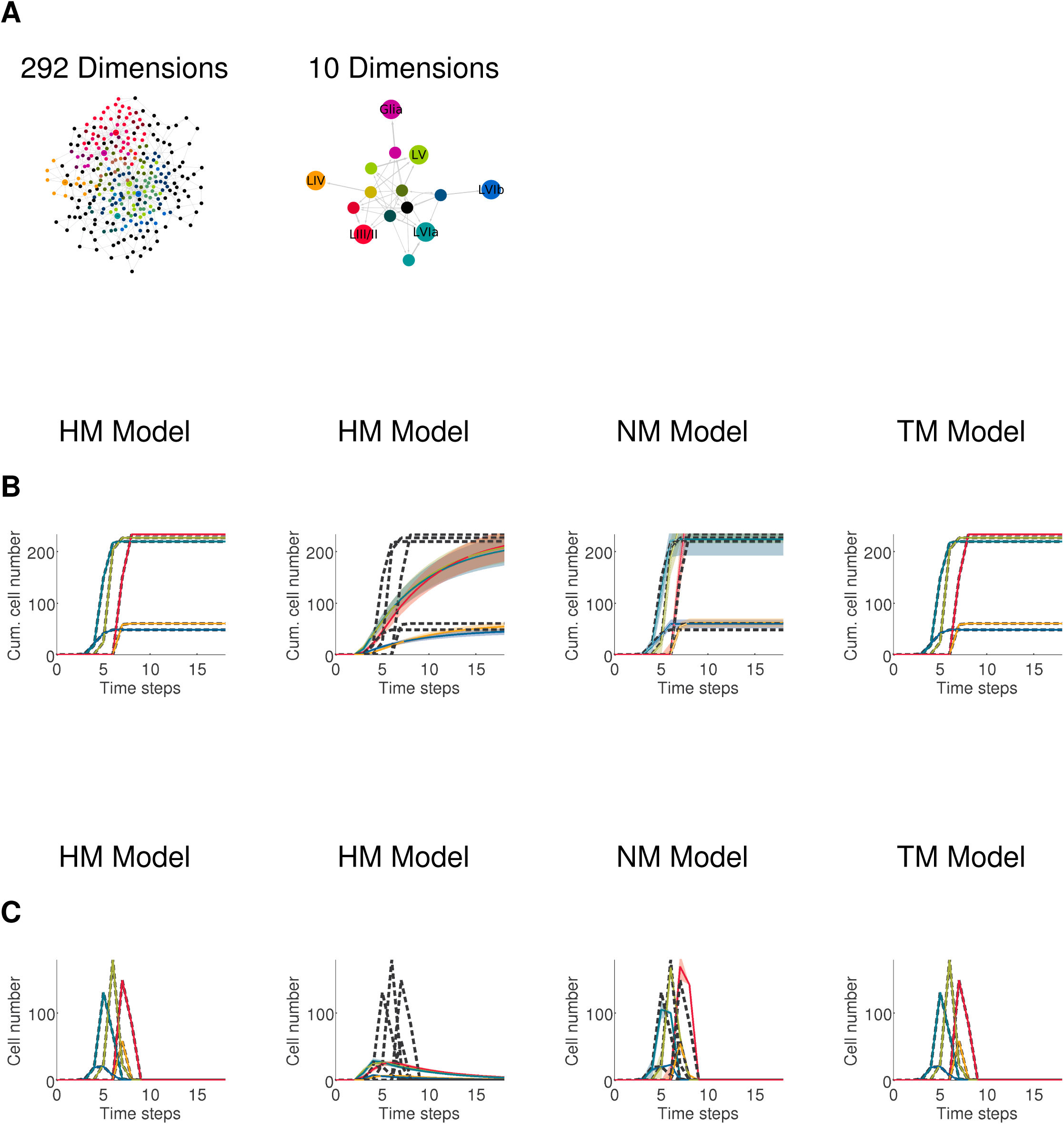
State Diagrams and model generated cell distributions for cortical area 6. (**A**) Original State Diagram *D* = 292 and its reduced *D* = 10 version for cell lineages in cortical area 6. Nodes represent cell states, arrows state transition probabilities. Cell state colors are the same as for Figure S5. (**B**) Generation of cells by various stochastic models. Mean cumulative number of differentiated cells produced at each time step. (**C**) Mean instantaneous number of differentiated cells produced at each time step. Dashed lines, original distribution; colored lines, model distribution; shaded area, standard deviation. HM, Homogeneous Markov model; NM, Non-homogeneous Markov model; TM, Time-dependent Markov model. Low-dimensional HM model fails to capture the data, whereas TM performs well.

**Figure S8.**
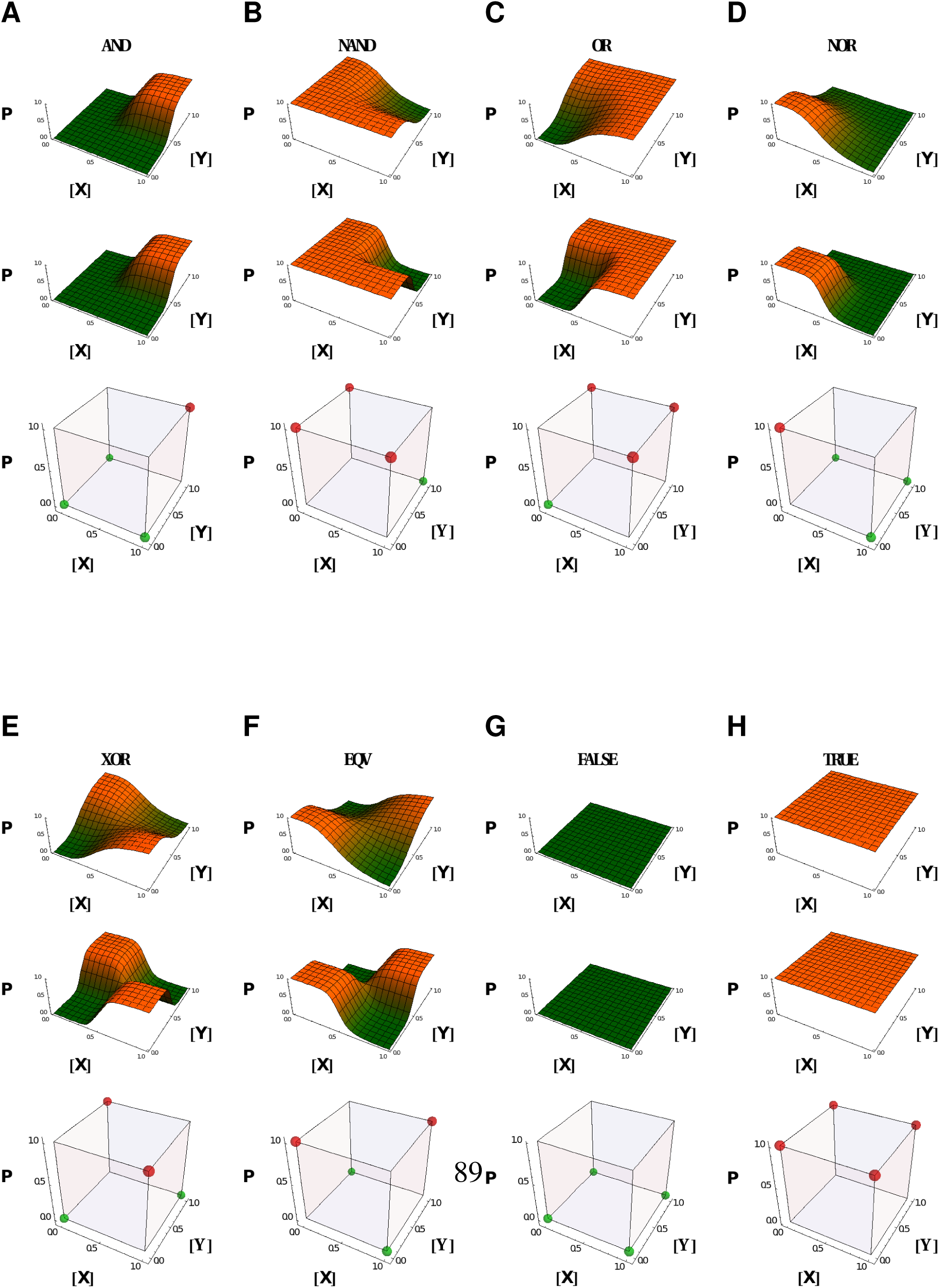
Combinatorial transcription logic. Cis-regulatory constructs can implement conventional canalizing logic gates (**A**) AND, (**B**) NAND, (**C**) OR, (**D**) NOR and non-canalizing (**E**) XOR, (**F**) EQV, (**G**) FALSE, (**H**) TRUE. The z-axis represents the output partition function *P* given [*X*] and [*Y*]. The computation depends on the steepness of the sigmoidal function *H*, ranging from (top to bottom row) continuous, approximately Boolean and discrete Boolean.

**Figure S9.**
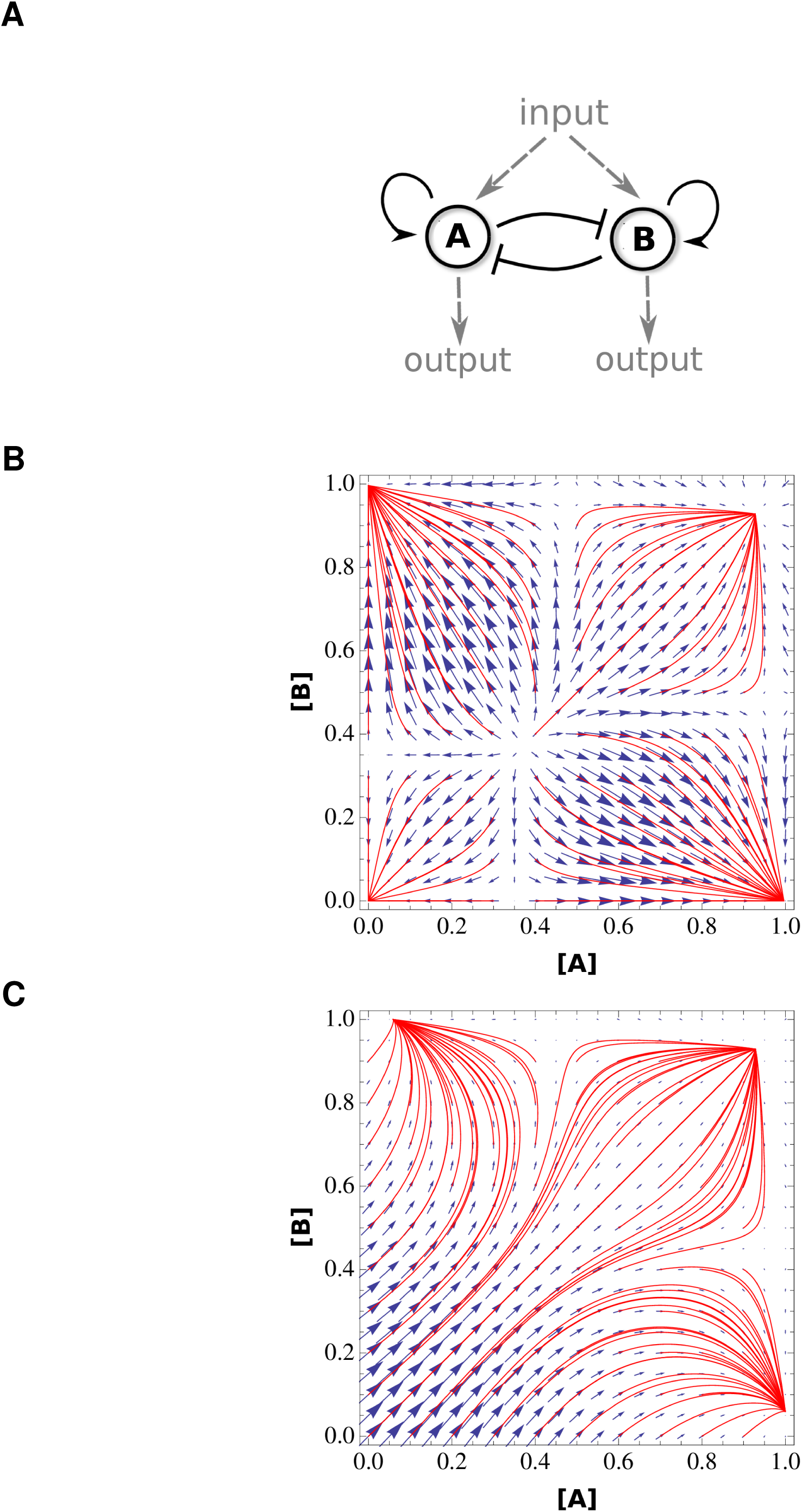
Dynamics of a 2-dimensional genetic switch. (**A**) Scheme of subnetwork with mutual inhibition between two transcription factors *A* and *B*, each with positive feedback; an external input *I*; and two outputs. (**B**) Vector field representing the gradient direction as a function of concentrations *A* and *B*, for switch without input (*I* = 0). The system has 4 attractor states, which means that the attractor states at high concentrations have hysteresis. (**C**) Vector field representing the gradient direction as a function of *A* and *B* for switch with input *I* = 1. Attractors at either high *A* or *B* represent downstream differentiation pathways. Red traces are simulated trajectories from various initial points.

